# Within and cross species predictions of plant specialized metabolism genes using transfer learning

**DOI:** 10.1101/2020.01.13.112102

**Authors:** Bethany M. Moore, Peipei Wang, Pengxiang Fan, Aaron Lee, Bryan Leong, Yann-Ru Lou, Craig A. Schenck, Koichi Sugimoto, Robert Last, Melissa D. Lehti-Shiu, Cornelius S. Barry, Shin-Han Shiu

## Abstract

Plant specialized metabolites mediate interactions between plants and the environment and have significant agronomical/pharmaceutical value. Most genes involved in specialized metabolism (SM) are unknown because of the large number of metabolites and the challenge in differentiating SM genes from general metabolism (GM) genes. Plant models like *Arabidopsis thaliana* have extensive, experimentally derived annotations, whereas many non-model species do not. Here we employed a machine learning strategy, transfer learning, where knowledge from *A. thaliana* is transferred to predict gene functions in cultivated tomato with fewer experimentally annotated genes. The first tomato SM/GM prediction model using only tomato data performs well (F-measure=0.74, compared with 0.5 for random and 1.0 for perfect predictions), but from manually curating 88 SM/GM genes, we found many mis-predicted entries were likely mis-annotated. When the SM/GM prediction models built with *A. thaliana* data were used to filter out genes where the *A. thaliana-*based model predictions disagreed with tomato annotations, the new tomato model trained with filtered data improved significantly (F-measure=0.92). Our study demonstrates that SM/GM genes can be better predicted by leveraging cross-species information. Additionally, our findings provide an example for transfer learning in genomics where knowledge can be transferred from an information-rich species to an information-poor one.

## Background

As more genome sequences become available, a major challenge in biology is to connect genotype to phenotype [1]. At the molecular level, phenotypes can be defined as products derived from genomic sequences, including transcripts, proteins, and/or metabolites. Plants produce a diverse array of specialized metabolites, with estimates upwards of 200,000 structurally unique compounds [2,3], many of which are important in medicine, nutrition, and agriculture [4–6]. Plant metabolic activities are broadly classified into two categories. The first is general (or primary) metabolism (GM), which involves the production of metabolites essential for survival, growth, and development in most, if not all, plant species [3,7]. In contrast, specialized (or secondary) metabolism (SM) leads to the accumulation of lineage-specific metabolites that may confer a fitness advantage in particular environments [2,3,8,9]. For example, some plant specialized metabolites such as glucosinolates and terpenoids confer resistance against insects and pathogens [6,10]. Another difference between general and specialized metabolites is that the later tend to accumulate in specific tissues such as in trichomes or fruit [11,12]. In addition to their ecological and evolutionary importance, specialized metabolites are important for human health; ∼25% of medicinal compounds are derived from plant metabolites [5,13]. For example, *Solanum nigrum* and *S. lyratum*, produce glycosides that have anti-tumor activity in cancer cell lines [14]. *Atropa belladonna*, nicknamed ‘beautiful woman’ because in Roman times women used its extract to dilate their pupils [15], is a producer of the tropane alkaloids hyoscyamine and scopolamine, has anticholinergic activity, and is used to treat parasympathetic nervous system disorders and asthma [16,17]. Furthermore, specialized metabolites contribute to desirable agronomic traits such as the aromas and flavors of fruits [11] and defense against agricultural pests [18].

Tomato is a model crop that has emerged as a system for investigating SM pathways. For example, the production of acylsugars, a specialized metabolite, in tomato and its wild relatives is important for repelling herbivores [19–21]. Some specialized metabolites found in the tomato fruit also confer health benefits by, for example, reducing risk of cancers and coronary heart diseases [4,22,23]. Despite recent progress in elucidating tomato SM pathways, our understanding of many of the steps in these pathways are incomplete due to the diversity of specialized metabolites. Many genes that underlie the production of specialized metabolites belong to the same gene families as genes involved in GM [8,24–26], which makes them difficult to distinguish. Currently, genetic approaches are used to identify SM genes in tomato, including gene silencing [27], genetic mapping [28], and the use of introgression lines [29]. In addition, genes involved in SM or belonging to a particular pathway can be predicted computationally. For example, protein sequence information can be used to predict enzymatic functions and assign genes to pathways [30–32], which can have high error rates [33]. Gene co-expression networks have also been used to classify genes into specific metabolic pathways [34]. In addition, involvement of genes in a pathway can also be hypothesized using correlation of gene expression with the production of specific metabolites [35–37]. Finally, heterogenous gene features including gene duplication status, evolutionary properties, expression levels, placement in co-expression networks, and protein domain content have been integrated using supervised machine learning to make SM/GM gene predictions in Arabidopsis [38].

Supervised learning approaches leverage instances (genes in this study) with known labels (SM or GM) to learn how the properties (i.e., features) of those instances can be best used to distinguish instances with different labels in the form of a predictive model (**Figure 1**). There are two factors limiting computational predictions of SM/GM genes. First, although supervised learning methods for SM/GM prediction are effective in Arabidopsis, it remains unclear how these methods may work in species with less complete gene and pathway annotations. Second, as sequence similarity-based approaches have high error rates, it is challenging to transfer annotation information across species [39]. The goal of this study is to address these limitations using an approach called “transfer learning” [40], where knowledge of SM/GM annotations from Arabidopsis was transferred to for predicting tomato SM/GM genes.

**Figure 1:**
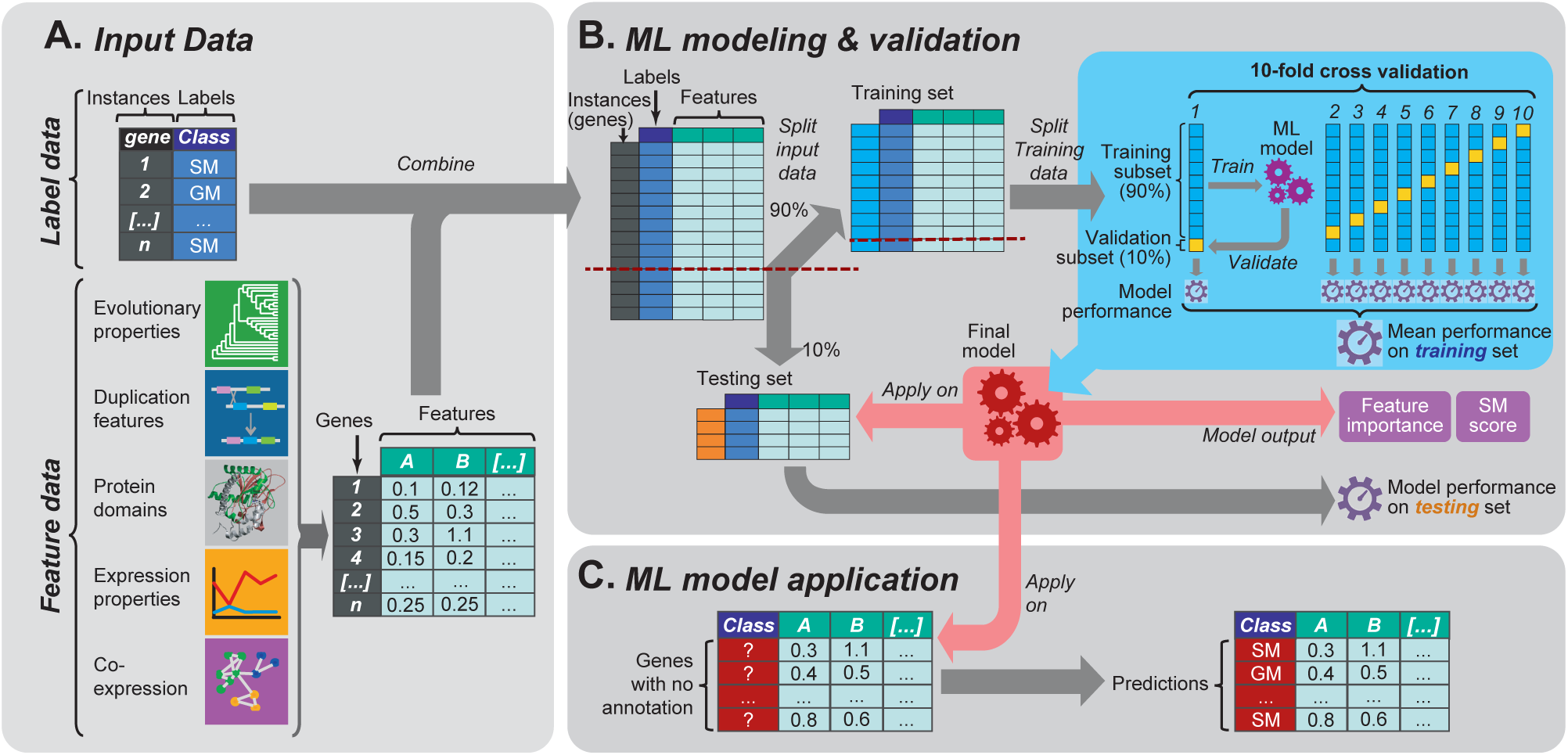
Machine learning workflow used in this study. **(A)** Schematic showing the input data for machine learning. The first inputs are labeled instances, collectively referred to as the model training set. In this case the instances are genes and the labels are the gene classes (response variable; either specialized or general metabolism, SM or GM). The second input is features, or the predictive variables in the model. In this study, five feature categories, which each contain multiple features, were utilized: evolutionary properties, duplication features, protein domains, expression properties, and co-expression data. Each gene (instance) has a value for each feature. **(B)** The machine learning process. First the data set was split into training (90%) and testing (10%) sets. Next, equal numbers of training instances (i.e., 500 GM and 500 SM genes) were randomly selected from the training set to learn prediction models. This step was repeated 100 times, with different subsets of GM/SM genes selected from the training set in each repeat, to assess the robustness of prediction models. For each repeat, a 10-fold cross-validation was performed where the selected instances were further divided into a training subset (90%) for building the model and a cross-validation subset (10%; distinct from the testing set withheld from model building) to evaluate the model. After cross-validation, the optimal parameters were chosen to establish the final model for a given training/feature data set. Model performance assessed using the cross-validation sets was represented using the average F-measure of all repetitions. In addition to assessing performance based on cross-validation, another F-measure was calculated for the final model based on its application to the testing set that was held out from the very beginning and never used for training. (**C)** The final model is applied on unannotated enzymatic genes to make predictions.

## Results and Discussion

### Identifying specialized metabolism genes in tomato using machine learning approaches

Prior to applying the transfer learning approach, we first used a supervised learning approach to build a model capable of classifying a tomato gene as either an SM or GM gene to serve as the “baseline” model for comparing against transfer learning results later on. For model training data, we used TomatoCyc annotated metabolic enzyme genes (referred to as “annotated genes”, see **Methods**, for annotation information see **Table S1**), where genes in pathways under the category “secondary metabolism biosynthesis” were considered SM genes (538 genes). Genes in any other pathway not under the SM category were considered to be GM genes (2,313 genes). Genes found in both SM and GM pathways (158) were excluded. The remaining annotated genes were divided into a training set (90%) for model training and a testing set (10%) for model performance evaluation. For all annotated tomato SM and GM genes (2,861), we collected and processed five gene feature categories (**Figure 1A**): evolutionary properties, gene duplication mechanism, protein domain content, expression values, and co-expression patterns (7,286 total features, see **Methods**, for feature values see **Dataset S1**). The values of these features for genes in the training set were then used to train multiple machine learning models for predicting whether a gene was likely an SM or GM gene (see **Methods, Figure 2A**).

**Figure 2:**
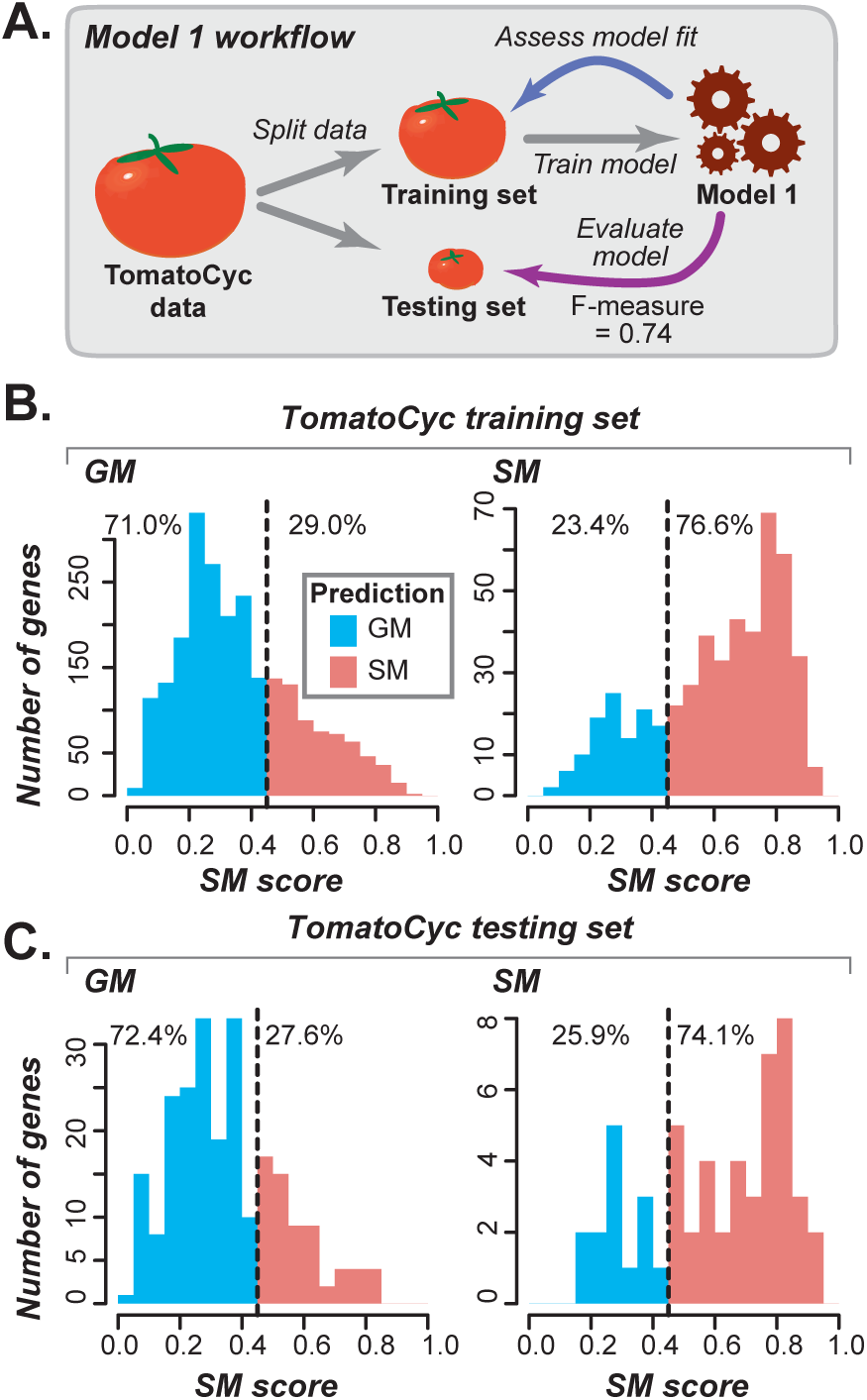
Tomato-based Model 1 and its performance. **(A)** Schematic illustrating Model 1, in which a tomato data set with 7,286 tomato features were used. The model was built using TomatoCyc annotations and applied to tomato genes. **(B)** Distribution of Model 1 SM gene likelihoods (SM scores) for the TomatoCyc-annotated SM and GM genes in the training set. Prediction threshold, based on the score with the highest F-measure, is indicated by the dotted line, and predicted GM (blue) and SM (red) genes are to the left and to the right of the line, respectively. Percentage values indicate the percent total genes predicted as GM or SM. **(C)** Distribution of Model 1 SM scores for testing SM and GM genes that were withheld from model training.

We determined model performance by calculating F-measure (the hormonic mean of precision and recall, see **Methods**). For other measure of model performance, see **Table S2**. The best performing model (Model 1) has F-measure = 0.74 (**Figure S1A**). The Model 1 F-measure is significantly better than a random guess (0.5) but far from perfect (1). Using Model 1, 76.6% of annotated SM genes and 71.0% of annotated GM genes had predictions consistent with their TomatoCyc annotations (**Figure 2B)**. To provide an independent validation, the model was then applied to the testing set, which resulted in a similar F-measure of 0.73 (**Figure 2C, Table S2**). Because the test set was withheld from model training, this indicated the model could be applied to genes with no annotation and provide reasonable predictions. By applying Model 1, each gene was given a likelihood score, referred to as the SM score (see **Methods**), which indicates how likely a particular gene is to be an SM gene (**Figure 2B**). For SM scores and SM/GM predictions for all tomato enzymatic genes for all models, see **Table S3**.

We identified features with the top 50 importance scores from Model 1 (**Figure S1B**, for feature importance for each model, see **Table S4**). The higher the importance score, the better the feature is at separating SM and GM genes. By and large, the important features for the tomato Model 1 is similar to those for predicting Arabidopsis SM/GM genes [38]. For example, similar to SM genes in Arabidopsis, tomato SM genes tend to be in larger gene families (median = 8) compared with GM genes (median = 3, **Figure 3A**; for test statistics, see **Table S5**), are more likely to be tandem duplicates (37%) than GM genes (13%), have a lower proportion as syntenic duplicates (17%) compared with GM genes (25%, **Figure 3B**), and have higher synonymous/synonymous substitution rates (*dN/dS*) relative to GM genes in both cross-species (**Figure 3C, Figure S2A-H**) or within species (**Figure 3D, Table S4**) comparisons. The lower the *dN/dS* value, the stronger the negative selective pressure a gene has experienced. Thus, SM genes were experiencing less intense negative selection compared to GM genes. We also found that many more homologs of tomato SM genes exist within species or in closely related species compared to GM genes (**Figure 3E**).

**Figure 3:**
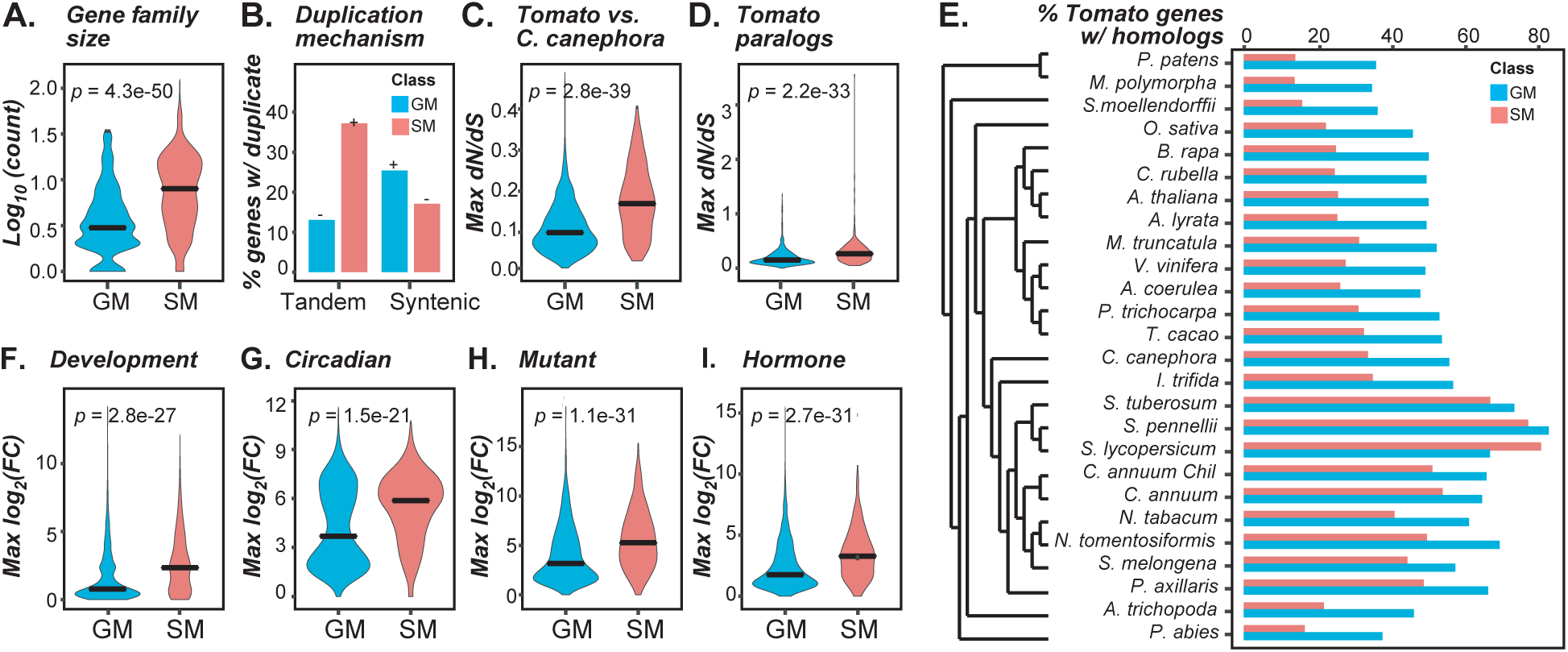
Duplication, evolutionary, and expression features important for Model 1 predictions of SM and GM genes. **(A)** Log 10 of gene family size (number of paralogs) for the families GM (blue) and SM genes belong to. All *p*-values are from the Mann-Whitney U tests between SM and GM genes. **(B)** Percent of GM and SM genes with at least one duplicates derived from tandem or syntenic mechanism. **(C)** Maximum *dN/dS* values from comparisons of tomato SM and GM genes to homologs in *C. canephora*. **(D)** Maximum *dN/dS* values of tomato SM and GM genes to their paralogs. **(E)** Phylogenetic tree of 26 species and a bar plot showing the percentage of tomato GM and SM genes that have at least one homologs in each species. **(F-I)** Distribution of maximum log 2 fold change (FC) between all samples in a given dataset for GM and SM genes in each class (GM and SM) over four datasets: **(F)** meristem development (1 study, 18 samples), **(G)** circadian (1 study, 86 samples), **(H)** mutant (14 studies, 239 samples, see **Table S6** for list of mutants) and **(I)** hormone treatment (5 studies, 89 comparisons, see **Table S6** for hormone treatments).

Variation in transcriptional levels and patterns between genes may represent differences in their functions and can therefore also be key features distinguishing SM and GM genes. We compiled 47 transcriptome studies (for details on the datasets, see **Table S6**) spanning a range of environmental conditions, hormone treatments, and developmental stages, mostly in wild-type genetic backgrounds. In Model 1, 147 out of the top 200 most informative features were related to expression (**Table S4**). For example, maximum log fold change between developmental stages, circadian time points, mutants vs. wild type, and hormone treatments vs. controls are among the top expression features (ranked 12-30, **Figure S1B, Table S4**). SM genes tended to have higher maximum fold change values (**Figure 3F-I, Table S5, S6**), but lower expression levels (**Figure S2I-J**) than GM genes. Thus, SM gene expression tends to be more variable across developmental stages, times of day, and environment. Consistent with this, expression variation (median absolute deviation, see **Methods**) is also an important feature (**Table S4**). For example, many specialized metabolites important for fruit flavor and color are produced during tomato fruit development [11]. Aside from gene expression, the enrichment of specific protein domains such as the p450 domain among SM genes (**Figure S2K**) is an additional feature that differentiates them from GM genes.

### Characteristics of genes with inconsistent annotations and predictions

Although the tomato SM/GM prediction model F-measure (0.74) was significantly better than a random guess (0.5), 29% of GM genes were mis-predicted as SM and 23% of SM genes were mis-predicted as GM when using an SM score threshold determined based on the optimal F-measure (**Figure 2B**). In addition, the tomato model did not perform as well as an earlier model for predicting Arabidopsis SM/GM genes (F-measure = 0.79, Moore et al., 2019). Note that the tomato model is trained on TomatoCyc annotations, which can be of poorer quality than those of AraCyc (Arabidopsis annotations)—there are only 16 experimentally verified TomatoCyc SM/GM genes compared to 1,652 in AraCyc. To understand why we obtained a high rate of mispredictions, we assessed what features may cause a gene to be mis-predicted. For example, SM genes in general tend to be in larger gene families than GM genes, and genes annotated as GM but predicted as SM (annotated→predicted: GM→SM) tended to belong to larger gene families (median = 5) than those having consistent GM annotations/predictions (GM→GM, median = 3, **Figure 4A**). Similarly, annotated SM genes predicted as GM (SM→GM) belonged to smaller families (median = 3) compared with correctly annotated/predicted SM genes (SM→SM, median = 10, **Figure 4A**). Additionally, we found that GM→SM genes tended to be tandem duplicates, similar to SM→SM genes and in contrast to GM→GM and SM→GM genes (**Figure 4B**). These findings indicate that mis-predicted genes tend to possess feature values that are deviated from the norms.

**Figure 4:**
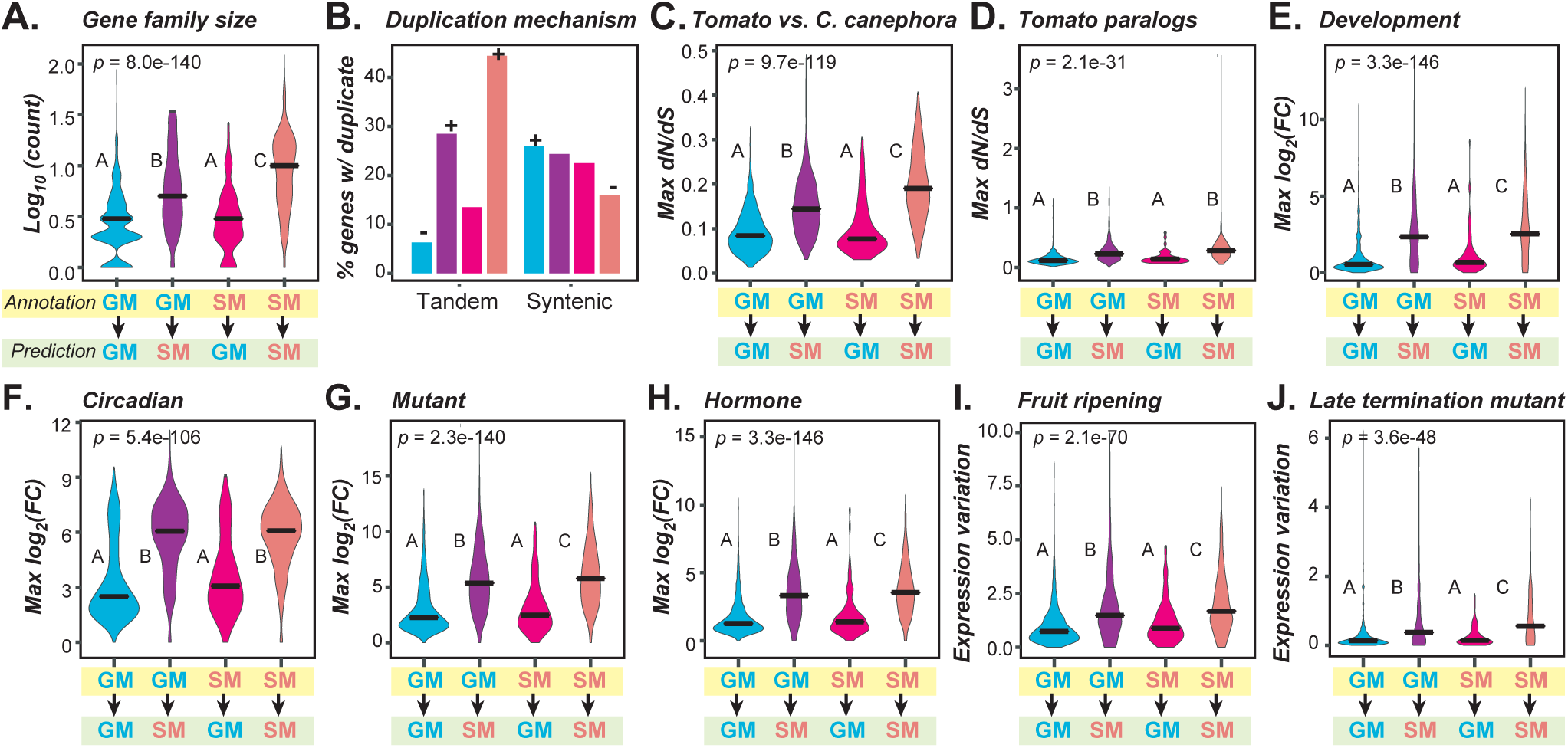
Feature distributions of genes with predictions contrary to their annotated classification. **(A)** Log 10 of gene family sizes among four Annotation (yellow rectangle)→Prediction (green rectangle) classes. Blue: GM→GM, a GM gene predicted as GM. Purple: GM→SM: a GM gene predicted as SM. Magenta: SM→GM: an SM gene predicted as GM. Red: SM→SM: an SM gene predicted as SM. For this and subsequent figure depicting continuous data, *p*-values are from the Kruskal-Wallis test and post-hoc comparisons were made using the Dunn’s test. Different letters indicate statistically significant differences between groups (*P* < 0.05). **(B)** Percentage of genes with at least one duplicates derived from tandem or syntenic mechanism. Color scheme following that in **(A)**. + and -: significant enrichment of SM and GM genes, respectively at 5% significance level after Benjamin-Hochberg multiple testing correction. **(C)** Maximum *dN/dS* values from comparisons of genes in four classes to homologs in *C. canephora*. **(D)** Maximum *dN/dS* values of genes in four classes to their paralogs. **(E-H)** Distributions of maximum fold change over the same expression data set as in **Figure 3F-I** including **(E)** the meristem development, **(F)** the circadian, **(G)** mutant, and **(H)** hormone treatments. **(I-J)** Distribution of fold change variation in two datasets: **(I)** fruit ripening (1 study, 12 samples) and **(J)** *late termination mutant* (1 study, 12 samples).

Another example where GM→SM and SM→GM genes defied the general trend was in maximum *dN/dS* value, having higher and lower *dN/dS* values, respectively, compared with those genes with consistent annotations/predictions (**Figure 4C,D, Figure S3A-H**). For example, one of the GM→SM genes, *XP_010323708* (*Solyc07g054880*.*3*.*1*), has a maximum *dN/dS* of 0.25 relative to its *Coffea canephora* homolog, which is much higher than that observed for GM→GM genes (*dN/dS* of 0.10) (**Dataset S1, Table S5**). This high *dN/dS* value likely contributed to the prediction of this gene as SM. When looking more closely at *XP_010323708*, we found that this gene was previously reported to encode a methylketone synthase that produces specialized methyl ketones specific to the *Solanum* genus [41], and should be annotated as an SM gene. Other GM genes with high *dN/dS* values from comparisons to their tomato paralogs were also predicted as SM genes. For example, three *Glycoalkaloid metabolism* (*GAME*) genes involved in steroidal glycoalkaloids production – *GAME4, GAME12*, and *GAME17* – stand out as SM genes in our model while TomatoCyc incorrectly annotated them as GM genes. *GAME4* and *GAME12* both have high maximum *dN/dS* values relative to tomato paralogs (0.30 and 0.26, respectively), a feature that many other SM genes share (SM median = 0.27, GM median= 0.15). *GAME17* belongs to a large protein family (30), another feature common to SM genes (SM median = 8, GM median = 3) and the most important feature for Model 1. In contrast to GM→SM genes, SM→GM genes have a maximum *dN/dS* score (median = 0.27) from comparisons to tomato paralogs that is significantly below that for SM→SM genes (median= 0.33, **Figure 4C, Table S5**). Aside from evolutionary properties and duplication features, compared with SM→SM genes, GM→SM genes also had similar maximum expression fold differences (**Figure 4E-H**), expression variation values (**Figure 4I, J**), median expression levels (**Figure S3I, J**), and protein domain compositions (**Figure S3K**).

In summary, we found that the distributions of feature values for mis-predicted GM→SM genes mirrored those for annotated SM genes. Likewise, the feature values distributions for SM→GM genes were similar to the overall distributions for annotated GM genes. These observations indicated that some SM genes in TomatoCyc looked more like GM genes and some GM genes looked more like SM genes which contributed to the discrepancies between annotation and prediction. An open question is whether these mis-predicted genes were misannotated in the first place or if they were correctly annotated but incorrectly predicted by a faulty model. This prompted us to look more closely at mis-predicted genes to see if their annotations were supported by compelling experimental evidence.

### Manual curation of SM/GM genes to obtain a benchmark set

Based on comparison of feature value distributions, mis-predicted genes tend to possess properties more similar to the class (GM or SM) they were mis-predicted as. This is not a surprising outcome because our explicit goal was to learn about generalizable differences between annotated GM and SM genes. The unresolved question is why mis-predictions occur. Three factors may account for mis-predictions: (1) the genes were annotated correctly, and Model 1 was incorrect, (2) Model 1 made correct predictions, but the annotations were incorrect, and (3) both annotations and predictions were correct, because these genes have roles in both GM and SM, i.e., they have dual functions (DF). To assess these possibilities, we manually curated a set of 88 tomato genes (83 with annotations in TomatoCyc) encoding enzymes classified as SM, GM, or DF based on published evidence of *in vitro* enzyme activity and/or *in planta* characterization (see **Methods**). These 88 genes are collectively referred to as the benchmark set, and the curated evidence supporting their SM/GM/DF designations are shown in **Table S1**.

Out of 31 TomatoCyc-annotated GM genes analyzed, 24, 5 and 2 were manually curated as GM, SM and DF genes, respectively. Among the five annotated GM genes that were manually curated as SM, all five were predicted by Model 1 as SM. Four are the aforementioned genes *Methylketone synthase* (*XP_010323708*), *GAME4, GAME12* and *GAME17*. The three *GAME* genes contribute to glycoalkaloid biosynthesis in several Solanaceae species [27]. The fifth gene correctly predicted by Model 1 is the neofunctionalized gene *Isopropylmalate synthase 3* (*IPMS3*), which acquired a role in an SM pathway after the duplication of an ancestral *IPMS* gene involved in amino acid metabolism (GM pathway). *IPMS3* is a tissue-specific SM gene involved in acylsugar production in glandular-trichome tip cells and is curated as an SM gene based on empirical evidence [42]. Thus, in these cases, Model 1 made the correct predictions, but the annotations were incorrect. Two *Geranylgeranyl diphosphate synthases (GGPS, NP_001234087* and *NP_001234302*) are manually curated as DF genes, but annotated by TomatoCyc as GM and predicted by Model 1 as SM. The challenge in classifying these genes might arise from the fact that GGPS enzymes catalyze core reactions in isoprenoid biosynthesis, an ancient and diverse pathway that leads to the synthesis of both GMs and lineage-restricted SMs [43].

Manual curation of 45 TomatoCyc-annotated SM genes revealed that 3 were likely GM genes and 5 were likely DF genes. We chose to look in detail at the three manually curated GM genes that were annotated as SM: two carotenoid biosynthesis genes, *PHYTOENE DESATURASE* and *TANGERINE* [44,45], and a cytochrome P450, *SlKLUH*, that, when mutated, disrupts chloroplast homeostasis and has pleiotropic effects on plant growth and development [46]. As carotenoid biosynthesis is conserved among all photosynthetic organisms [47], and disruptions in basic development processes, such as gametophyte and seed development, is an indicator of essentiality in all plants [48], these genes should be considered GM genes. In all three cases, Model 1 predictions agreed with the TomatoCyc SM annotations and, thus both the predictions and annotations were incorrect.

Next, we focused on comparing the manually curated benchmark set to Model 1 predictions. We found that 17 out of 29 (58.6%) total benchmark GM genes, and 13 of the 24 benchmark GM genes that were annotated as GM by TomatoCyc (54%), were incorrectly predicted as SM by Model 1 (**Figure S4A**; **Table S3**). Thus, Model 1 tended to mis-predict benchmark GM genes as SM genes. In contrast, of the 51 total benchmark SM genes, 45 (88.2%) were correctly predicted by Model 1 (**Figure S4A**; **Table S3**). Taken together, our Model 1 predictions were mostly consistent with the SM benchmark classifications. However, the model clearly had trouble predicting known GM genes. With regard to TomatoCyc-annotated genes, the opposite was true – 24 of 29 (82.8%) benchmark GM genes were correctly annotated as GM, and 37 of 47 (78.7%) benchmark SM genes were correctly annotated as SM. Therefore, for SM gene prediction, Model 1 has a lower error rate (11.8%) compared with the TomatoCyc annotation (21.3%), indicating that a higher proportion of benchmark SM genes were annotated in TomatoCyc than GM genes. However, for benchmark GM genes, Model 1 has a higher error rate (46% of benchmark GM genes predicted as SM genes) than the TomatoCyc annotation (14.3% of benchmark GM genes predicted as SM).

### Using transfer learning to make predictions across species

Based on analysis of the benchmark data, there are two major sources for mis-predictions. The first is that a subset of the TomatoCyc-annotated SM or GM genes were incorrectly annotated, and these mis-annotations were propagated into Model 1. The second is that Model 1 predict these genes correctly. These two explanations are not mutually exclusive, and the extent to which each contributes to mis-predictions remains to be determined. To determine the most likely reason for the mis-predictions and to improve upon Model 1, we used both the benchmark gene set and the TomatoCyc annotations to build a new model (referred to as Model 2), but this did not improve the prediction accuracy (F-measure=0.74, same as Model 1, **Figure S1A, Table S2**). This was likely due to the small proportion of benchmark gene-inspired annotation corrections (30) relative to the large number of TomatoCyc-annotated genes (2,858).

We next asked whether information from Arabidopsis, which diverged from the tomato lineage 83-123 million years ago [49,50], could be used to improve gene predictions in tomato. Here we use a machine learning approach called transfer learning [40] in which a base model is first built using data from Arabidopsis and then the learned features and/or the base model itself are used to make predictions in tomato using the tomato annotations and features. To accomplish this, a list of 4,197 similar features in Arabidopsis and tomato (referred to as shared features, see **Methods**) were identified. A model was built using previously defined AraCyc GM/SM annotations [38] and shared features. This model is referred to as Model 3 (**Figure 5A**) that performed reasonably well in separating *A. thaliana* GM/SM genes (**Figure 5B**). For comparison, we also built a model (Model 4) using TomatoCyc GM/SM annotations and tomato data for the same shared features as in Model 3 and to train the model (**Figure 5C**). Model 3 built with Arabidopsis shared feature data had an F-measure = 0.81 when it was used to predict Arabidopsis genes as GM/SM (**Table S2**). In comparison, Model 4 built with tomato shared feature data had an F-measure = 0.75 when used for predicting tomato annotations (**Table S2**). Additionally, more GM/SM genes in Arabidopsis are predicted correctly by Model 3 (**Figure 5B**) than GM/SM genes in tomato by Model 4 (**Figure 5D**).The higher F-measure and better predictions for Model 3 are consistent with there being more experimentally based gene annotations for Arabidopsis than for tomato that likely contribute to the differences in model performance.

**Figure 5:**
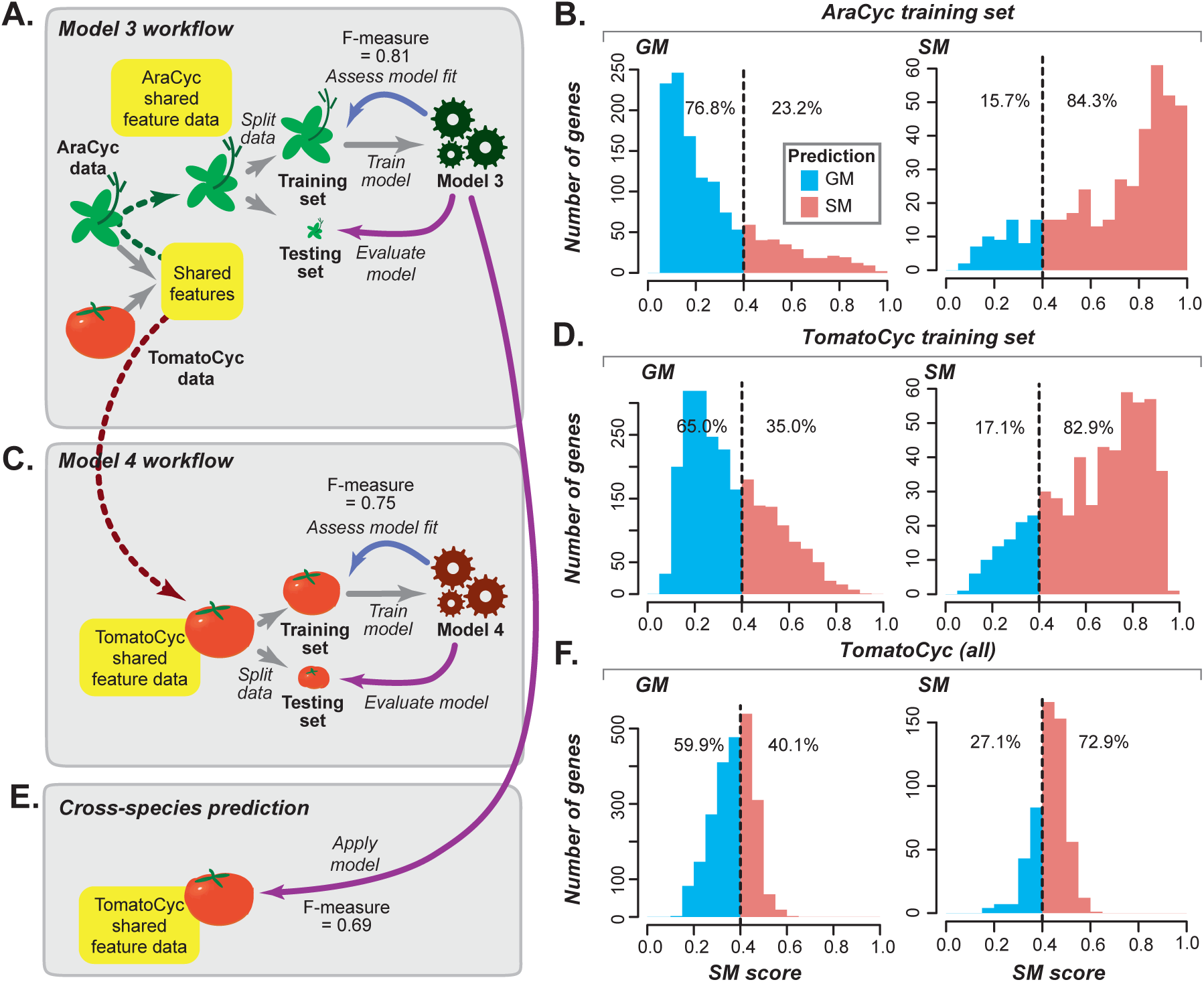
Arabidopsis Model 3 and tomato Model 4 predictions. **(A)** Model 3 built with the Arabidopsis training data using only shared feature set between Arabidopsis and tomato. **(B)** Distribution of Model 3 SM scores of Arabidopsis SM and GM training set genes. Dotted line: same as in **Figure 2**. Blue and red: SM and GM genes, respectively. **(B)** Model 4 built with the tomato training data using only shared feature set between Arabidopsis and tomato. **(D)** Distribution of Model 4 SM scores of tomato training set GM and SM genes. **(E)** Application of Model 3 on tomato genes using the shared feature set between Arabidopsis and tomato. **(F)** Distribution of Model 3 SM scores of tomato genes.

We next applied Arabidopsis-based Model 3 to predict tomato SM and GM genes and obtained an F-measure of 0.69 (**Figure 5E, Table S2**). This was substantially lower than the F-measure obtained when applying tomato-based Model 4 to tomato genes (0.75, **Table S2**), and fewer TomatoCyc annotated GM/SM genes were predicted correctly (**Figure 5F**). Based on SM scores for these models, 21.1% of TomatoCyc GM genes were predicted as GM genes by tomato Model 4 but predicted as SM genes by Arabidopsis Model 3 (lower right quadrant, **Figure 6A, Table S3**). However, Model 3 predicted 50% of benchmark tomato GM genes as GM (**Figure S4B**), which – although far from perfect – is substantially better compared with the percentage of benchmark GM genes correctly predicted by tomato Model 4 (25%, **Figure S4C**). Thus, Arabidopsis data (when used to train Model 3) led to improved tomato GM gene predictions compared with tomato annotation data. Based on our finding that annotated GM genes were more likely to be misannotated compared with annotated SM genes (**Figure S4B, C**), this indicates that the decline in model performance was due to mis-annotation of tomato genes.

**Figure 6:**
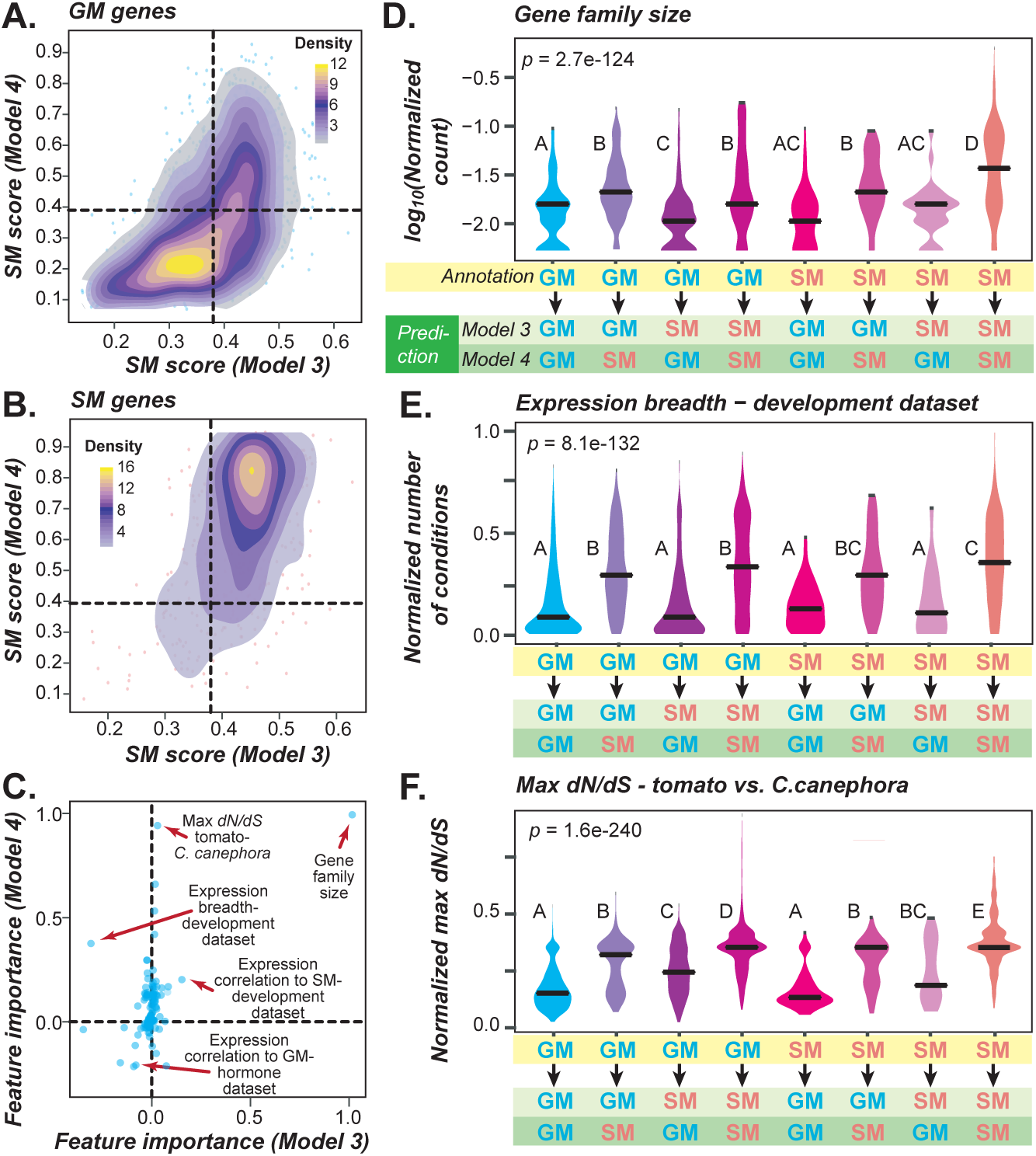
Tomato Model 4 and Arabidopsis Model 3 comparison. **(A)** Correlations between TomatoCyc GM genes SM scores based on Model 3 and Model 4. Color: data point density ranges from high (yellow) to medium (purple), to low (fading purple). **(B)** Correlation of TomatoCyc SM genes SM scores based on Model 3 and Model 4. **(C)** Correlations in feature importance values based on Model 3 and Model 4. Arrows point to example consistent and inconsistent features. **(D-F)** Feature value distributions for annotated SM and GM genes that are predicted as SM or GM genes by Model 3 and Model 4. *P*-values are from Kruskal-Wallis tests and post-hoc comparisons were made using the Dunn’s test. Different letters indicate statistically significant differences between groups (*P* < 0.05). **(D)** Log 2 of normalized gene family size. **(E)** Normalized expression breadth based on the meristem development data. **(F)** Normalized maximum *dN/dS* between tomato genes and their homologs in *C. canephora*.

### Reasons why Arabidopsis-based Model 3 had suboptimal performance on tomato genes

To further assess the possibility of mis-annotation, we asked how well Model 3 and 4 predict benchmark SM genes. We found that benchmark tomato SM genes were less well predicted using Arabidopsis Model 3 (84% correctly predicted, **Figure S4B**), a substantial drop from the near perfect predictions (97%) using tomato Model 4 (**Figure S4C)**. This indicated that Arabidopsis data may provide more useful information about true GM genes in other species than about SM genes, likely because GM genes are conserved among plant species, and many have been studied using Arabidopsis as a model. Thus, it is more straightforward to transfer knowledge about Arabidopsis GM genes to tomato. SM genes, in contrast, are by definition lineage-specific and not all SM gene properties will be shared across species, which explains the drop in prediction accuracy in Model 3 compared with Model 4. Nonetheless, the SM likelihood scores are largely consistent between Models 3 and 4 (**Figure 6A, B**; **Figure S5A, B**; **Table S3**), indicating there remain substantial similarities among SM genes across species.

When we looked into the models in more detail, we found that the major reason why Arabidopsis Model 3 predicted genes differently from tomato Model 4 is because they have different important features (**Figure 6C**). Aside from the three most consistently important ones, which are gene family size, expression correlation between SM genes during development, and expression correlation between GM genes in the hormone dataset (**Figure 6C**), many features such as maximum *dN/dS* relative to *C. canephora* homologs are highly important in tomato Model 4 but much less important in Arabidopsis Model 3. Upon examination of feature value distributions, we found that, in general, the feature values of the tomato Model 4-based predictions more closely aligned with those of the annotated genes in the tomato training set than with Arabidopsis Model 3-based predictions (**Figure 6D-F**). For example, annotated tomato SM genes predicted as GM genes by Arabidopsis Model 3 but as SM genes by tomato Model 4 (referred to as SM→GM_3_/SM_4_ genes, the plot in pink, **Figure 6D**) tend to be in large gene families like SM→SM_3_/SM_4_ genes (the orange plot, **Figure 6D**). In contrast SM→SM_3_/GM_4_ genes (the brown plot, **Figure 6D**), tend to be in small gene families. This indicates that tomato Model 4 is more strongly influenced by gene family sizes when differentiating SM and GM genes than Arabidopsis Model 3. This general pattern is also true for expression-based and *dN/dS* features (**Figure 6E, F**; **Figure S5C-F**). For example, GM→GM_3_/SM_4_ genes are likely predicted as SM genes by tomato Model 4 (the second plot, **Figure 6F**) because they have high *dN/dS* values similar to those of the SM genes used to train the model (the eighth plot, **Figure 6F**). However, GM→SM_3_/GM_4_ genes (the third plot, **Figure 6F**) tend to have lower *dN/dS* values similar to those of the GM genes used to train the model (the first plot, **Figure 6F**). In the above example, the Arabidopsis Model 3 yields predictions contrasting with those from tomato Model 4. Most notably, the Arabidopsis Model 3-based predictions have feature values that mostly defy the general trends of the GM and SM genes in the tomato training data. This indicates that there are differences between the training data for Arabidopsis Model 3 and tomato Model 4 that bias each model.

### Improving the tomato-based model by removing potentially mis-annotated genes identified based on the Arabidopsis model predictions

We hypothesized that if the Arabidopsis Model 3-based predictions are correct, then the genes with contrasting predictions and annotations are mis-annotated and their removal from the training data would lead to significantly improved predictions. This is because training the model from incorrect examples (i.e., mis-annotated entries) will lead to suboptimal models making erroneous predictions. On the other hand, if the Arabidopsis Model 3-based predictions are completely uninformative, the removal of genes from the training set would not improve the prediction. Thus, to further test the above hypotheses, we removed TomatoCyc-annotated GM and SM genes that had contradictory predictions from Arabidopsis-based Model 3 (i.e. GM→SM_3_ and SM→GM_3_) from the training set. Using this filtered training data set, a new tomato data-based model, Model 5, was generated using the same shared feature set between Arabidopsis and tomato for Model 3 and 4 (**Figure 7A**, see **Methods**).

**Figure 7:**
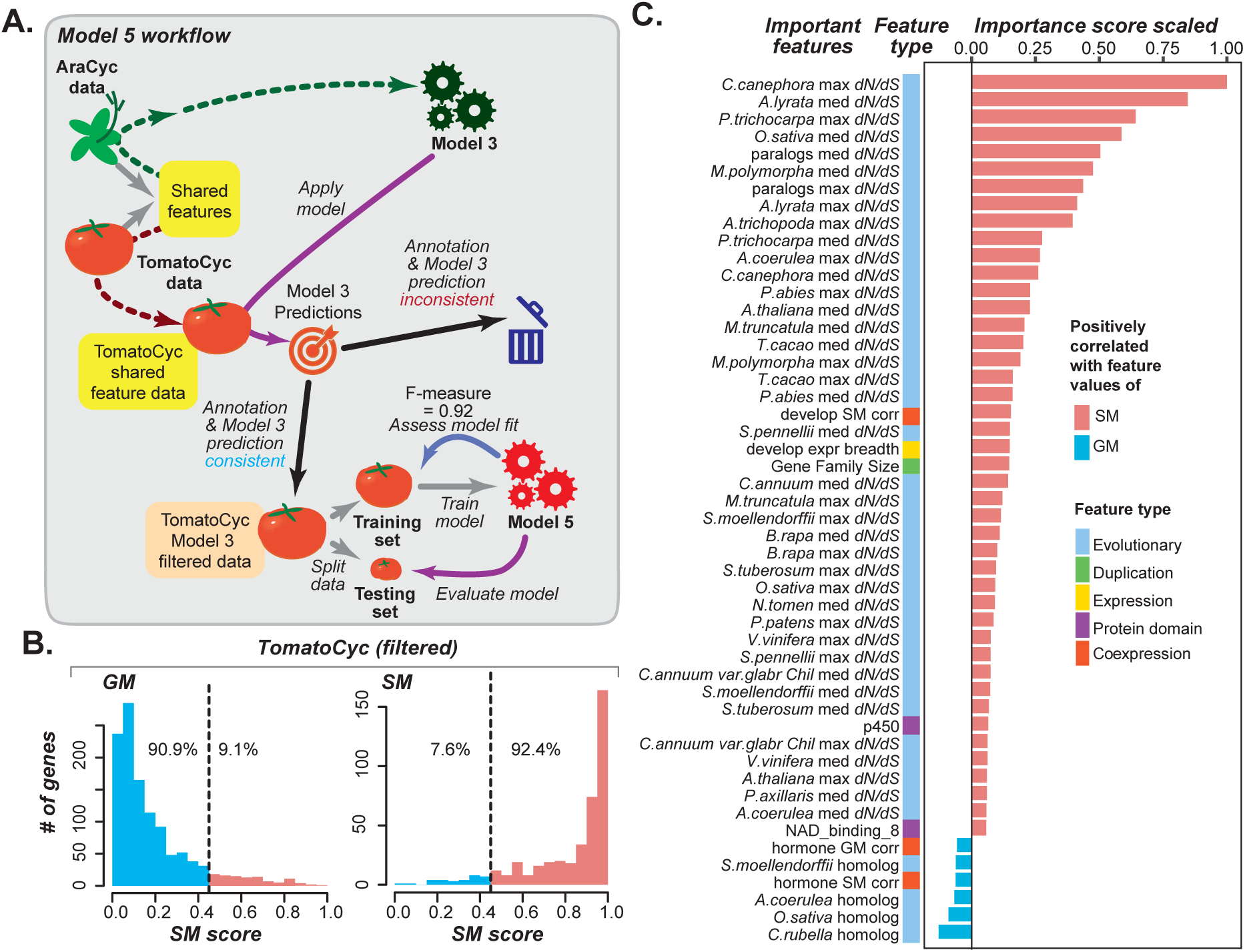
Model 5 performance and important features. **(A)** Diagram showing the procedures leading to Model 5 for predicting tomato GM/SM genes. **(B)** Distribution of Model 5 SM scores of tomato training set GM and SM genes. Dotted line: same as in **Figure 2**. Blue and red: SM and GM genes, respectively. **(C)** Feature importance values for Model 5. Blue: importance scores normalized to between -1 and 0 for top features positively correlated with GM gene feature values (more negative is more important). Red: importance scores normalized between 0 and 1 for top features positively correlation with SM gene feature values (more positive is more important).

When we applied this filter to build tomato Model 5, there was a dramatic improvement in tomato GM/SM gene predictions (F-measure = 0.92, **Figure S1A, Table S2**) compared with predictions based on Model 3 (F-measure= 0.69, **Figure S1A, Table S2**) and Model 4 (F-measure = 0.75, **Figure S1A, Table S2**). In particular, we were able to predict 90.9% of all annotated GM genes and 92.4% of all annotated SM genes in the filtered training data as GM and SM genes, respectively (**Figure 7B, Table S2**). Thus, Model 5, trained on a data set where GM→SM_3_ and SM→GM_3_ genes have been removed, is significantly improved compared with previous models. To validate Model 5 with an independent dataset, we applied it to a testing set of 159 SM and GM genes withheld from Model 5 during training. We found that 84% and 88% of the test set GM and SM genes, respectively, were predicted consistently with their annotations (**Figure S6B**).

To test whether model improvement was due to the filtering out of a subset of misannotated genes from the tomato training data and not just to the removal of genes in general, we built 10 additional models (collectively referred to as Model 6) using the same number of tomato SM and GM training genes as used for training Model 5, except that the genes were removed randomly. We found the median F-measure to be the same as that from Model 4 (where no SM or GM genes were removed; **Figure S1A, Table S2, see Methods**), showing no model improvement. Thus, the improvement in model performance of tomato Model 5 could not be attributed to random gene removal and was likely achieved because the filtered tomato training data did not contain mis-annotated genes that would confuse the model.

After showing that Model 5 performed significantly better on training data, we next asked how Model 5 faired in predicting benchmark GM genes. We found that 75% of benchmark GM genes were correctly predicted by Model 5 (**Figure S6A, Table S3**), compared with 25% for tomato Model 4 and 50% for Arabidopsis Model 3 (**Figure S4F, G**). In contrast, there was no improvement in benchmark SM predictions when comparing Model 4 (94% correct, **Figure S4F, Table S3**) to Model 5 (92% correct, **Figure S6A, Table S3**). These findings indicate that the improvement in Model 5 is likely due to its ability to determine true GM genes while maintaining true SM gene prediction performance. In addition, our results suggest that the filtering step mostly corrected for GM genes misannotated as SM genes in TomatoCyc. Consistent with this conclusion, 83.1% of the annotated SM genes that were removed from the Model 5 training data because Model 3 called them as GM, were predicted as GM genes by Model 5 (**Figure S6C**). This indicates that introducing GM genes that were likely misannotated as SM genes into the training set led to a sub-optimal model. After their removal, the new model was able to better identify GM genes misannotated as SM. In contrast, among annotated GM genes removed from the training set because they were predicted as SM genes by Model 3, only 6.1% were predicted by Model 5 as SM genes (**Figure S6C**). Furthermore, GM genes identified as SM genes by Model 3, were mostly still predicted as GM genes, indicating that the removal of these genes was relatively inconsequential, and the main issue was that a substantial number of GM genes were mis-annotated as SM genes.

Additional models (Models 7 and 8) were trained using the same filtered gene set used in training Model 5 but with the full tomato feature data set (instead of just the shared features used in Models 3, 4, and 5; **Figure S6D**). The training set for Model 8 also included the benchmark gene annotations. Models 7 and 8 had similar performances (F-measure = 0.88 and 0.86 respectively, **Table S2, Figure S6E-G**). Both Models 7 and 8 were significantly improved compared with Model 1 (F-measure = 0.74), particularly when predicting GM genes (similar to Model 5). Overall, using Arabidopsis Model 3 to remove potentially mis-annotated tomato genes, i.e. genes that were not good training examples, led to substantially improved models (Model 5 and 7), especially for predicting GM genes.

While TomatoCyc provides annotations for many genes in SM pathways, the global SM gene content in tomato is unknown. To provide a genome-wide estimate of SM gene content in the tomato genome, we used Model 7 to classify 5,627 unannotated enzyme genes and found that 2,865 are likely involved in SM pathways (**Figure S6H**). This indicates that substantially more SM genes are yet to be identified because only 696 genes are currently annotated in TomatoCyc. As noted earlier, each enzyme gene has an SM score from the model application, which can be interpreted as the probability that a gene is an SM gene (see **Table S3** for scores for each gene); thus, those unannotated enzymes that are highly likely to be an SM gene can be prioritized for further investigation.

### Relationships between improved performance and feature rankings

Models 5 and 7 substantially improved gene predictions in tomato compared with all other models because mis-annotated genes, mostly genes annotated as SM but predicted as GM by Arabidopsis Model 3, were removed from the training data. To better understand the reasons for the improvement in GM gene predictions, we looked into three examples where Models 5 and 7 predicted manually curated GM benchmark genes as GM genes, but where tomato-based Models 1 and 4 predicted the genes as SM genes: *1-aminocyclopropane-1-carboxylate oxidase 1* (*LeACO1*, NP_001234024), *abscisic acid 8’-hydroxylase* (*CYP707A1*, NP_001234517), and the cytochrome P450 *SlKLUH* (XP_004236064). In these cases, the mispredictions were likely due to gene expression-related features. While *LeACO1* exhibited a maximum log_2_ fold change of 7.0 based on the fruit ripening dataset (**Dataset S1**), which is consistent with the higher values observed for SM genes (median=1.9) than for GM genes (1.2, *p=*1.3e-15). Similarly, the variance of log_2_ fold change in expression during fruit ripening for *SlKLUH* is 2.5, which is consistent with significantly higher median variance for SM genes (1.5) compared with GM genes (1.0, *p=*1.9e-21). *CYP707A1* is up-regulated under many developmental conditions (13), which is not typical for tomato GM genes (SM median =16, GM median = 9, *p*=9.3e-26). Additionally, the expression of *LeACO1, CYP707A1*, and *SlKLUH* correlates highly with that of other SM genes (PCC= 0.87, 0.63, and 0.83, respectively). The similarity of these expression feature values as those of SM genes likely contributed to their mis-prediction by Models 1 and 4.

Importantly, Models 5 and 7 likely predict these three genes correctly as GM genes because of the reduced reliance of these models on features associated with gene expression. Models 1 and 7 both use the full feature set, but filtered training data were used to train Model 7. In Model 1, expression variance in fruit ripening was ranked 46 among important features, while in Model 7 it was ranked 120 (**Table S4**). Similarly, when comparing Models 4 and 5, which both use the shared feature set but differ in whether filtered training data were used, the features expression breadth under development and expression correlation between SM genes were ranked higher for Model 4 (6 and 16, respectively) than for Model 5 (22 and 20, respectively) (**Table S4**). Model improvement is also due to higher ranking of evolutionary features, such as maximum *dN/dS* between tomato genes and *C. canephora* homologs, median *dN/dS* between tomato genes and homologs in *Arabidopsis lyrata*, and maximum *dN/dS* between tomato genes and homologs in *Populus trichocarpa*. In Model 5 these features were ranked 1, 2, and 3, respectively; in Model 4 they were ranked 2, 3, and 8, respectively; **Table S4**); in Model 7 they were ranked 1, 2, and 7, respectively; and in Model 1 they were ranked 2, 9, and 16, respectively **Table S4**. *LeACO1* and *CYP707A1* both have maximum *dN/dS* values from comparisons to *C. canephora* homologs (0.07) more similar to those of GM genes (median=0.10) than to SM genes (0.17). Similarly, *SlKLUH* has a maximum *dN/dS* value from comparisons to *A. lyrata* of 0.11, which is closer to the GM median (0.09) than to the SM median (0.15). Because in Models 5 and 7 these *dN/dS* features were weighted more heavily and certain expression features were weighted less heavily, the *dN/dS* feature values contributed to their correct classification as GM genes.

In addition to the features discussed thus far, we also found that gene family size was no longer the most important feature in Models 5 and 7, ranked 24 and 27, respectively, as it was Models 1, 3 and 4. Considering that some of the largest enzyme families - such as cytochrome P-450 and terpene synthases - contain both SM and GM genes, this reduced importance likely contributed to improved predictions. Despite the improvement, Models 5 and 7 are by no means perfect and erroneous predictions still occur. For example, *PSY1* is a fruit ripening-related gene manually curated as an SM benchmark gene, but it was predicted as a GM gene by both Models 4 and 5. *PSY1* represents an unusual case of duplication-associated sub-functionalization and is specifically expressed in chromoplast-containing tissues such as ripening fruits and petals [51]. *PSY1* has comparatively low *dN/dS* values (similar to GM genes), especially between tomato and *C. canephora* (maximum *dN/dS* = 0.06). Because this *dN/dS* feature was the most important feature for Model 5, this ultimately contributed to the misprediction of *PSY1* as a GM gene.

Other examples are two GM terpene synthases involved in the biosynthesis of gibberellin, a plant hormone [52]: *copalyl diphosphate synthase* (*CPS*, NP_001234008) and *kaurene synthase* (*KS*, XP_004243964). Both *CPS* and *KS* are mis-predicted as SM genes in all models, presumably because of their high *dN/dS* values from comparisons to homologs in several species (*CPS* median *dN/dS*= 0.20, *KS* median *dN/dS*= 0.26). These two enzymes were derived from an ancestral dual functional enzyme containing both copalyl diphosphate synthase and kaurene synthase activities [7]. Angiosperm terpene synthases seem to have lost one activity or the other, but the ancient timing of the CPS/KS duplication (after divergence between bryophytes and the other land plant lineages) makes the high rate of evolution unusual. It is unknown what effect the loss of activity has on the evolution of the terpene synthase sequence. For all three genes, *PSY1, CPS*, and *KS*, the atypical evolutionary rates, either unusually low or high, led to mis-prediction. Overall, our machine learning approach led to a highly accurate SM/GM model with an F-measure of 0.91 (where a value of 1 indicates a perfect model). However, while our approach ensures the identification of typical SM/GM genes, SM/GM genes with atypical properties that defy the general trend still are likely mis-predicted.

### Conclusions *# Conclusions section is not encouraged*

Many SM genes are unknown due to the vast number of specialized metabolites are limited to specific species and SM and GM genes are difficult to distinguish because SM genes are often derived from GM genes. Additionally, many specialized metabolites of interest are found in medicinal plants or crops that are not well annotated. If data from a better annotated species such as Arabidopsis could be used, directly or indirectly, to make cross-species predictions in another species, such as tomato, this could greatly improve annotations in non-model species. Here we used machine learning to establish models for classifying genes with SM and GM functions in tomato, but consistent with the lower quality of the tomato annotation, these models established using tomato features had relatively poor performance compared with models built in Arabidopsis. We also found that a substantial number of important features and predictions differed between the models based on Arabidopsis (Model 3) and tomato (Model 4). We discovered that the differences in feature importance and model performance were likely the result of mis-annotation of some tomato genes, which contributed negatively to the performance of machine learning models. Therefore we attempted to perform cross-species knowledge transfer by using a machine learning approach called transfer learning [40], where knowledge learned from a previously trained model (e.g., our Arabidopsis Model 3) is used (in this case, to remove predictions inconsistent with annotations) to train another model (e.g., tomato Model 5). By filtering out TomatoCyc-annotated genes that had predictions opposite from those of the Arabidopsis-based Model 3 from the training data, we significantly improved the accuracy of tomato SM/GM gene predictions. We demonstrated that this improvement would not have been possible without informed removal of potentially mis-annotated data. This approach can be applied more generally to any problem in a species that is relatively information poor by transferring knowledge from an information-rich one. Using transfer learning we may also be able to better annotate less well studied species.

It is important to note that a limitation of the transfer learning approach is that it is only useful for transferring knowledge, mechanisms, or phenomena that are similar across species. In our study, the transfer learning approach worked well for GM genes, but it did not have an appreciable impact on the prediction of SM genes, likely because SM pathways are by definition specialized–what you learn in one species does not necessarily apply to another. A specific example of where transfer learning can suffer is in predicting genes with atypical properties. The machine learning approach excels at spotting patterns in data, and the performance of machine learning models improves as more high-quality instances (e.g., experimentally validated SM/GM genes) and more informative features (e.g., *dN/dS*) are incorporated. However, it is a challenge to generate high-quality instances, and expert knowledge dictates what kinds of features are incorporated. In addition, the representation of genes that are considered “atypical” in the model can be limited by our ability to scour the literature for novel features to represent these genes.

In future studies, transfer learning can be used to predict GM genes and, to a lesser extent, SM genes in species that lack annotations and/or experimental evidence such as non-model, medicinal plant species. An open question in this area that needs to be addressed is whether more closely related species, even though they may not be as well annotated, are better candidates for transfer learning than better annotated but more distantly related species. In addition, as discussed above, our models can potentially be further improved by incorporating additional features, particularly those that are shared between species, using transfer learning. For example, data that are incorporated as features for across species models should come from experiments performed in more similar ways in terms of treatments applied and tissues investigated. Furthermore, we found that SM gene annotations can vary across species, so reliance on information from a particular species may skew the model predictions and the features that are most important for the model. Thus, in future studies comparisons between models using data from single and multiple species can potentially inform further efforts to improve cross-species predictions via transfer learning. Another consideration is that we treated our research problem as a binary (SM or GM) classification problem. Over the course of evolution, SM pathways may branch off from GM pathways or some SM pathways may ultimately become GM pathways because of increasingly wider taxonomic distribution. Thus, the extent to which a gene is considered to be SM is likely continuous, where genes at the end of an SM pathway may be more “SM-like” than genes at the beginning of the pathway, which may be linked to GM pathways. The question is how to define the degree of involvement of a gene in SM pathways and determine whether continuous SM scores, where GM and SM genes have low and high scores, respectively, are good proxies for involvement in these pathways. This can be accomplished by mapping SM scores to pathways to see if they are predictive of where a gene lies in a pathway.

## Methods

### Annotation

Only enzyme genes were included in this study. A gene was considered to be an enzyme gene if it had an EC or RXN number annotation in TomatoCyc or assigned using E2P2 v3.0 [31]. Tomato pathway annotations were downloaded from the Plant Metabolic Network Database, TomatoCyc v. 3.2 [32]. Pathways that were nested under “Secondary Metabolism Biosynthesis” or “Secondary Metabolites Degradation” were considered specialized metabolism (SM) pathways and genes within those pathways were considered SM genes. All other pathways were considered to be general metabolism (GM) pathways. If a gene was annotated as being in both an SM pathway and a GM pathway, the gene was considered to be dual function (DF). Additionally, the biosynthesis of plant hormones was considered GM even though some hormone pathways fell under the DF category. If a pathway was nested under both “secondary metabolism biosynthesis” and other general biosynthesis categories, the pathway was determined to be DF. For specific SM pathway annotations, the path ID from TomatoCyc was used.

### Benchmark genes

The benchmark gene set was identified based on expert knowledge and literature mining. Tomato genes were defined as GM, SM, or DF based on *in planta* functional analyses of mutant generated through gene silencing or knockout mutations and/or studies of *in vitro* biochemical activity. For the identity of the benchmark genes (i.e. manually curated as SM, GM, or DF genes), the evidence used for manual curation, and publications supporting the evidence, see **Table S1**.

### Features used for machine learning

All gene feature values can be found in **Dataset S1**. These 7,286 features are divided into several categories, each with different numbers of features: protein domains (4,232 features), expression value (280), co-expression (2,670), evolution (78), and gene duplication (26). Protein domain Hidden Markov Models from Pfam v.30 (pfam.xfam.org/) was used to identify protein domains in annotated tomato protein sequences with HMMER (https://www.ebi.ac.uk/Tools/hmmer/https://www.ebi.ac.uk/Tools/hmmer/) using the trusted cutoff, then a binary matrix for each gene and domain was created where 1 indicates the protein sequence of a gene has a given domain and 0 indicates it does not.

### Expression value features

For expression value features, RNA-seq Sequence Read Archive (SRA) files for tomato were downloaded from National Center for Biotechnology Information (NCBI; https://www.ncbi.nlm.nih.gov/) totaling 47 studies and 926 samples (**Table S6**). These data sets included development (13 studies including fruit, flower, leaf, trichome, anther, and meristem tissues), hormone-related (5 studies: cytokinin, auxin, abscisic acid, gibberellic acid, and auxin inhibitor treatments), mutant (14 studies which compared various mutants against wild type), stress treatment (16 studies including shade, various pathogens, cold, light, and heat treatments), and circadian (1 study with 60 samples). RNA-seq data were processed to determine both fold change and fragments per kilobase of transcript per million mapped reads (FPKM) (https://github.com/ShiuLab/RNAseq_pipeline). The SRA files were converted to fastq format and filtered with Trimmomatic [53] for sequence quality with default settings. Bowtie (http://bowtie-bio.sourceforge.net/bowtie2/index.shtml) was used to create the genome index from the tomato NCBI *S. lycopersicum* genome 2.5, then RNA-seq reads were mapped to the tomato genome using TopHat [54]. Samples with <70% mapped reads were discarded. Cufflinks was then used to obtain FPKM values for mapped reads [55]. HTSeq [56] was used to get raw counts for fold change analysis. Fold change analysis was performed using edgeR version 3.22.5 [57]. Using each data set individually or all data sets combined, the median and maximum, and variation values for each gene were calculated. For breadth of differential expression, the number of conditions under which a gene was up- and down-regulated was determined using log fold change values for each data set or combination of data sets. A gene was considered up-regulated if it had a log fold change > 1 and a multiple-testing corrected *p*-value < 0.05 and down-regulated if it had a log fold change < -1 and a corrected *p*-value < 0.05.

### Co-expression features

For co-expression features, expression correlation was calculated using three methods: Pearson’s Correlation Coefficient (PCC), Spearman’s correlation, and Partial Correlation (Corpcor). For each enzymatic gene (annotated and unknown), its expression correlation with each annotated SM/GM/DF gene was calculated (excluding self-correlation) using each method, each expression measure (fold change or FPKM) and each individual expression dataset (with a distinct Gene Expression Omnibus GSE number), combination of datasets, and all datasets combined (see **Table S6**). Then, for an enzymatic gene, E, the median and maximum of the correlation values of gene E for each class (SM, GM, or DF) of genes was determined and used as feature values. Next, tomato genes were clustered into co-expression modules using six methods (k-means, c-means, complete/average/ward hierarchical clustering, and weighted correlation network analysis) across each individual expression dataset, dataset combination, and all datasets combined (same as for expression correlation). This was done using both fold change and FPKM values. Using Random Forest from Python package Scikit-Learn [58], the top 200 co-expression modules that were the best for distinguishing SM and GM genes for each clustering method were selected to be part of the feature matrix for the models.

### Evolutionary features

Orthologs and duplication nodes were determined using OrthoFinder [59]. For input, protein sequence files from 26 different species were downloaded from Phytozome (https://phytozome.jgi.doe.gov/pz/portal.html), Sol Genomics Network (SGN, https://solgenomics.net/), PlantGenIE (http://plantgenie.org/), or NCBI (www.ncbi.nlm.nih.gov/genome): *Physcomitrella patens* 318 v3.3 (Phytozome), *Marchantia polymorpha* 320 v3.1 (Phytozome), *Selaginella moellendorffii* 91 v1.0 (Phytozome), *Picea abies* V1.0 (PlantGenIE), *Amborella trichopoda* 291 v1.0 (Phytozome), *Oryza sativa* 323 v7.0 (Phytozome), *Brassica rapa* 277 V1.3 (Phytozome), *Capsella rubella* 183 V1.0 (Phytozome), *Arabidopsis thaliana* 167 TAIR10 (Phytozome), *Arabidopsis lyrata* v2.1 (Phytozome), *Medicago truncatula* 285 Mt4.0v1 (Phytozome), *Vitis vinifera* 145 *Genoscope* 12x (Phytozome), *Aquilegia coerulea* V3.1(Phytozome), *Populus trichocarpa 210 v3*.*0* (Phytozome), *Theobroma cacao* 233 v1.1 (Phytozome), *Coffea canephora* (SGN), *Ipomoea trifida* V1.0 (NCBI), *Solanum tuberosum* V3.4 (SGN), *Solanum pennellii* SPENNV200 (NCBI), *Solanum lycopersicum* V2.5 (NCBI), *Capsicum annuum* CM334 v.1.55 (SGN), *Capsicum annuum var. glabriusculum* V2.0 (SGN), *Nicotiana tabacum* TN90 AYMY-SS NGS (SGN), *Nicotiana tomentosiformis* V01 (NCBI), *Solanum melongena* r2.5.1 (SGN), and *Petunia axillaris* V1.6.2 (SGN).

To identify putative orthologs, OrthoFinder was first run using default settings, including a BLAST run using protein sequence data for each pair of species with default parameters (E-value<0.001), markov clustering (inflation parameter=0.1) to create initial orthogroups, and dendroblast to create distance matrices between protein sequences of genes within each initial orthogroup. Initial gene trees were created using OrthoFinder. Three initial orthogroups were found to contain a single copy gene from each of the 26 species. Protein sequences of genes in each of these three orthogroups were aligned with MAFFT [60], and the alignment was used to build a phylogeny with RAXML (-m PROTGAMMAJTT -number of bootstraps 100 -outgroups Mpoly, Ppaten). This putative species tree was used as input into OrthoFinder to reconcile the gene trees for redefining orthogroups. Genes were considered to be homologous if they were in the same orthogroup. *dN/dS* (non-synonymous to the synonymous substitution rate ratio) was calculated with the yn00 program using PAML version 4.4.5 [61]. Gene family size was determined by the number of genes in an orthogroup within the species *S. lycopersicum*.

Duplication mechanism was determined using MCScanX-transposed [62]. Four duplication mechanisms were used as features: 1) syntenic duplicates: paralogous genes present in within-species collinear blocks; 2) dispersed (transposed) duplicates: for a pair of paralogs in species A, only one of their corresponding orthologs in species B is present in the inter-species syntenic block; 3) tandem duplicate: a gene is adjacent to its paralog; 4) proximal duplicates: a gene is separated by no more than 10 genes from its paralog. Genomic clustering features were derived from the genome annotation *Solanum lycopersicum* V2.5. A gene pair X and Y was considered to be in the same genomic cluster if gene X was located within 10 kbps downstream of the 3’-end or upstream of the 5’-end of gene Y, and X and Y were within 10 genes from each other. For gene X, the numbers of genes that qualified as Ys were determined separately for Ys in SM and GM pathways. The time point of the most recent duplication was determined from the most recent speciation node associated with each gene as determined by OrthoFinder [59]. Duplication nodes ranged from most ancient (Node 0) to most recent (Node 24). The most recent duplication points for genes appearing to originate from multiple duplication nodes were defined by the highest-numbered node they belonged to (**Figure S7**). Pseudogenes in tomato were determined as in Wang et al. (2018) where genomic regions with significant similarity to protein-coding genes but with premature stops/frameshifts and/or were truncated were treated as pseudogenes [63]. Detailed methods and parsing scripts for different features can be found in: https://github.com/ShiuLab/SM-gene_prediction_Slycopersicum.

### Statistics

Statistical calculations were performed using R and Python. For discrete features, their relationships with SM/GM designations were determined by the Fisher’s exact test. For continuous data, either the Mann Whitney U test (for comparing two groups) or the Kruskal-Wallis test followed by Dunn Pairwise Comparisons (for >2 groups) were used for tests of significance. Statistical results are in **Table S5**.

### Machine learning models

Multiple prediction models were made using the Python Sci-kit learn package [58] with two algorithms, Random Forest (RF) and Support Vector Machine (SVM). The pipeline (**Figure 1**) used to run the models can be found here: https://github.com/ShiuLab/ML-Pipelinehttps://github.com/ShiuLab/ML-Pipeline. For each model, 10% of the data was withheld from training as an independent, testing set. The remaining 90% was used for training. Because the dataset was unbalanced (2,321 GM genes, 537 SM genes), 100 balanced datasets were created from random draws of GM genes to match the number of SM genes. Using the training data, grid searches over the parameter space of RF and SVM were performed. The optimal hyperparameters identified from the search were used to conduct a 10-fold cross-validation run (90% of the training dataset used to build the model, the remaining 10% used for validation, **Figure 1**) for each of the 100 balanced datasets. In total eight models were established using different feature and training datasets as described in **Results & Discussion**. For a subset of models, feature selection using RF was implemented to reduce the features to 50, 100, 200, 300, 400, 500, and 1000 to determine the optimal number of features. Model performance was evaluated using F-measure, the harmonic mean of precision (proportion of predictions that are correct) and recall (proportion of genes correctly predicted). Each model outputs an SM score for each gene that is defined as the mean of predicted class probabilities of a sample to be in the SM class based on all decision trees in the forest. For each tree, the SM class probability was the fraction of genes predicted as SM. The threshold of the SM score used to determine if a gene was an SM or GM gene was the SM score value when the F-measure was maximized. The models also have an importance score for each input feature, which takes into account the weight of the feature by assessing how well the feature (node) splits the data between SM and GM genes in a decision tree in the “forest” and this is weighted by the proportion of samples reaching that node (impurity score). The decrease in impurity score from each decision tree is averaged across all decision trees in the forest so that the higher the number, the more important the feature [64,65].

### Shared features between Arabidopsis and tomato

**Dataset S2** lists the shared features and their values for Arabidopsis and tomato. For binary data, the features that were shared by both species were kept. These included two types of binary features: (1) protein domains: ∼4,000 Pfam domains common between Arabidopsis and tomato; (2) evolutionary features: presence of a homolog in one of the 26 species, pseudogene paralog, and tandem paralog, and whether the most recent duplication events took place in the lineages leading to the nodes shared by both species (nodes 0-7). The shared features also included the following continuous features: gene family size, genomic cluster gene count, median/maximum *dN/dS* values between genes and their homologs in each of the 26 species, median/maximum *dN/dS* values between genes and their paralogs, and expression-based features. To generate shared expression features, expression data were placed into four categories – abiotic, biotic, hormone, and development – in both species. For each category, the Arabidopsis expression breadth, breadth of differential expression, and co-expression correlation values using PCC were obtained from an earlier study [38]. The same sets of features were generated for tomato in this study. Continuous values were normalized within each species so that they would be comparable across species. For the normalization script see https://github.com/ShiuLab/SM-gene_prediction_Slycopersicum.

## Supplemental Figures

***Supplemental Figure 1: Comparison of all model scores and feature importance values for Model 1***

(A) Comparison of model scores. F-measure is shown on the y-axis and model is shown on the x-axis. Model type is denoted by color. Gray indicates Models 1-8 variants (i.e., different ML algorithms and/or numbers of features used) that are not described in the text. RF: Random Forest. SVM: Support Vector Machine. featsel25-1000: <u>feat</u>ures <u>sel</u>ected, sets of 25 to 1000. For model names, see **Table S2**. (B) Bar plot of the top 50 most important features for Model 1. The importance score is on the y-axis and all scores are normalized to the score of the most important feature, which was set as 1. Red bars represent features that are enriched for SM genes while the blue bars represent features enriched for GM genes. Features are listed along the x-axis, with the color denoting the feature category.

***Supplemental Figure 2: Important features for Model 1***.

(A-K) Distributions or bar plots of feature values for TomatoCyc-annotated SM and GM genes. (A-J) Significance determined by the Mann-Whitney U test. (A-H) Distributions of the maximum or median *dN/dS* value for a given gene relative to their homolog in *P. patens, S. moellendorffii, A. trichopoda, O. sativa, B. rapa, A. coerulea, P. trichocarpa* and *S. pennellii*. (I, J) Distributions of log 10 of median FPKM values for the Inflorescence data set and Root data set. (K) Percent of genes with a given Pfam domain. Overrepresentation (+) and underrepresentation (-) was determined using those genes with a *p*-value less than 0.05 from a Fisher’s Exact test between SM and GM genes with Benjamin-Hochberg multiple testing correction.

***Supplemental Figure 3: Features important for SM vs. GM predictions***

For all distributions of each predicted class, GM→GM represents GM genes predicted by Model 1 as GM, GM→SM represents GM genes predicted by Model 1 as SM, SM→GM represents SM genes predicted by Model 1 as GM, and SM→SM represents SM genes predicted by Model 1 as SM. Significant differences between continuous variables were determined by the Kruskal-Wallis test (A-J) and post-hoc comparisons were made using Dunn’s test. Different letters indicate statistically significant differences between groups (*P* < 0.05). For binary data (K), overrepresentation (+) and underrepresentation (-) were determined by the Fisher’s Exact test where (+) is significant overrepresentation of a predicted class and (-) is significant underrepresentation. A *p*-value < 0.05 after Benjamin-Hochberg multiple testing correction was considered significant. (A-H) Distributions of the maximum or median *dN/dS* value for a given gene from comparisons to its homolog in *P. patens, S. moellendorffii, A. trichopoda, O. sativa, B. rapa, A. coerulea, P. trichocarpa* and *S. pennellii*. (I, J) Distributions of log10 (median FPKM) values for the Inflorescence (I) and Root (J) data sets. (K) Percentage of genes with a given Pfam domain.

***Supplemental Figure 4: SM likelihood scores for manually annotated genes***

(A-D) Distribution of SM likelihood scores for manually annotated benchmark genes from Model 1. TomatoCyc-XX indicates the TomatoCyc annotation and BM-XX indicates the benchmark annotation. SM likelihood score is shown on the x-axis, number of genes is on the y-axis. Prediction threshold, based on the score with the highest F-measure, is indicated by the dotted line, and predicted SM genes are shown to the right of the line in red while predicted GM genes are shown to the left of the line in blue. (E) Bar plot showing the percentage of manually annotated benchmark genes predicted as SM or GM by Model 3. The original annotation from TomatoCyc is shown first, followed by the benchmark annotation and then the prediction. (F) Same as (E), except that the predictions were made using the tomato Random Forest (RF) model (Model 4) with shared features.

***Supplemental Figure 5: S. lycopersicum and A. thaliana model comparison and model performance***

(A-B) Comparison of the SM score distributions for tomato Model 4 (y-axis) and Arabidopsis Model 3 (x-axis). Support Vector Machine (SVM) and a shared feature set were used for both models. Density of data points ranges from high (yellow) to medium (blue-purple) to low (white). (A) SM scores for GM genes; (B) SM scores for SM genes; (C-F) Feature distributions for annotated SM and GM genes that are predicted as SM or GM genes by Arabidopsis Model 3 and tomato Model 4. The x-axis lists the annotations for each group of genes predicted using Arabidopsis Model 3 and tomato Model 4. *P*-values are from the Kruskal-Wallis test and post-hoc comparisons were made using the Dunn’s test. Different letters indicate statistically significant differences between groups (*P* < 0.05). (C) maximum Pearson’s Correlation Coefficient (PCC) between a given gene and all other SM genes under stress conditions; (D) maximum PCC between a given gene and all other SM genes during development; (E) maximum PCC between a given gene and all other GM genes under hormone treatment; (F) normalized median *dN/dS* values between tomato or Arabidopsis genes and their homologs in *O. sativa*.

***Supplemental Figure 6: Benchmark and test set predictions from finalized models with A. thaliana mis-predictions removed (Models 5 and 7)***

(A) Bar plot showing the percentage of manually annotated benchmark genes predicted as SM or GM by Model 5. The original annotation from TomatoCyc is shown first, followed by the benchmark annotation and then the prediction. Distributions of SM likelihood scores are shown in plots B, C, G, and H. (B) Model 5 test set SM and GM genes, which were held out from the model building process completely. (C) TomatoCyc SM and GM genes with annotations opposite to Arabidopsis Model 3 predictions removed from the filtered training set. (D) Schematic diagram showing the application of tomato Model 7 to tomato. The full tomato feature dataset was used to build a binary model using TomatoCyc SM and GM annotations after removing genes mis-predicted by Arabidopsis Model 3. The model was then applied to tomato genes. (E) TomatoCyc filtered training set SM and GM genes from tomato Model 7. (F) Bar plot showing the percentage of manually annotated benchmark genes predicted as SM or GM by Model 7. The original annotation from TomatoCyc is shown first, followed by the benchmark annotation and then the prediction. (G) Model 7 test set: SM and GM genes, which were held out completely from the tomato Model 7 building process and (H) unannotated tomato enzymes. For plots (B, D, F, and G): SM likelihood score is shown on the x-axis, number of genes is on the y-axis. Prediction threshold, based on the score with the highest F-measure, is indicated by the dotted line, and predicted SM genes are shown to the right of the line in red while predicted GM genes are shown to the left of the line in blue.

***Supplemental Figure 7: Speciation nodes***

Phylogenetic tree of 26 species showing speciation nodes (N0-N24). Most recent gene duplication node in text refers to the speciation node where gene was last duplicated.

## Supplemental Tables

***Table S1: Tomato gene annotation information***

Annotation information based on TomatoCyc and manual annotation.

***Table S2: Model scores***

Scores and information for all models.

***Table S3: SM gene scores***

SM prediction scores for all genes for each of the models.

***Table S4: Feature Importance***

Feature importance scores for all models discussed in the text.

***Table S5: Feature Statistics***

Statistics for original and shared features.

***Table S6: Transcriptome studies***

Information about all expression datasets used in the models.

***Dataset S1: Original features***

Dataset includes all of the features used for Models 1, 2, 7, and 8.

***Dataset S2: Shared features***

Dataset includes all of the shared features between Arabidopsis and tomato used for Models 3, 4, and 5.

## Declarations

### Ethics approval and consent to participate

Ethics approval was not applicable.

### Consent for publication

Consent for publication was not applicable

### Availability of data and materials

Data and materials availability are as noted in the method sections.

### Competing interests

The authors declare that they have no competing interests.

### Funding

This work was supported by a postdoctoral fellowship from the National Science Foundation (NSF) IOS-1811055 to C.A.S; NSF grant IOS-1546617 to R.L., C.S.B, and S.-H.S.; U.S. Department of Energy Great Lakes Bioenergy Research Center (BER DE-SC0018409) grant to R.L. and S.-H.S.; Michigan AgBioResearch and U.S. Department of Agriculture National Institute of Food and Agriculture Hatch project number MICL02552 to C.S.B; and NSF grant DEB-1655386 to S.-H.S.

### Authors’ contributions

BMM and SHS conceived the study; all authors contributed to the design of the study; BMM, PW, and AL implemented the computational analysis; PF, BL, YRL CAS, KS, and CSB contributed to manual annotation; BMM, PW, AL, and MDL contributed to interpretation of computational analysis results, all authors contributed to interpreting annotation results; all authors contributed to the manuscript draft.

## Acknowledgements

We thank the Shiu lab, Last lab, and Barry lab members for their help during the preparation of this manuscript.

## Notes

#### Summary of Updates

Updated figures and texts.

