## Supplementary material for "Within and cross species predictions of plant specialized metabolism genes using transfer learning": Figure S1-7

#### Supplemental Figure 1

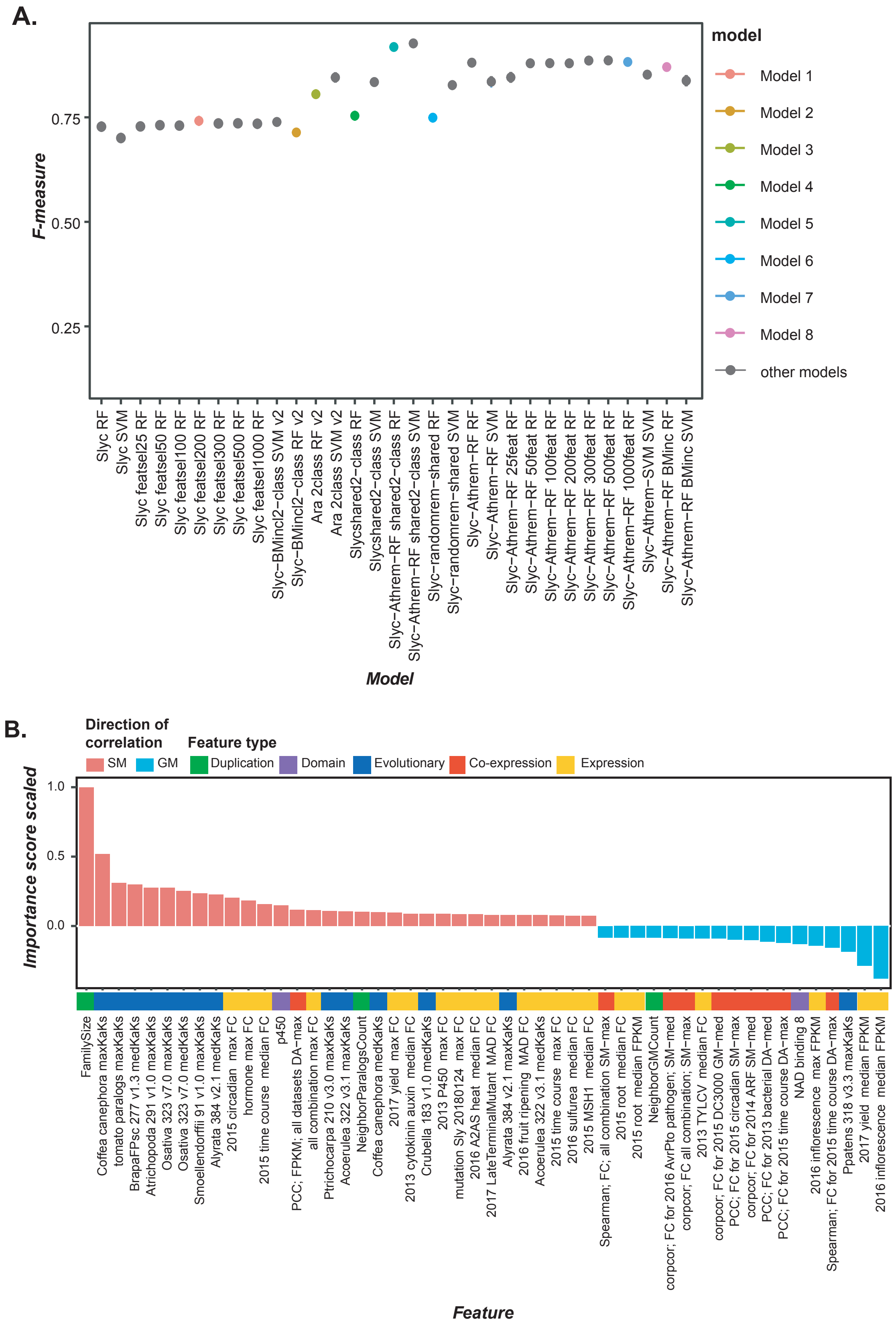

#### Supplemental Figure 2

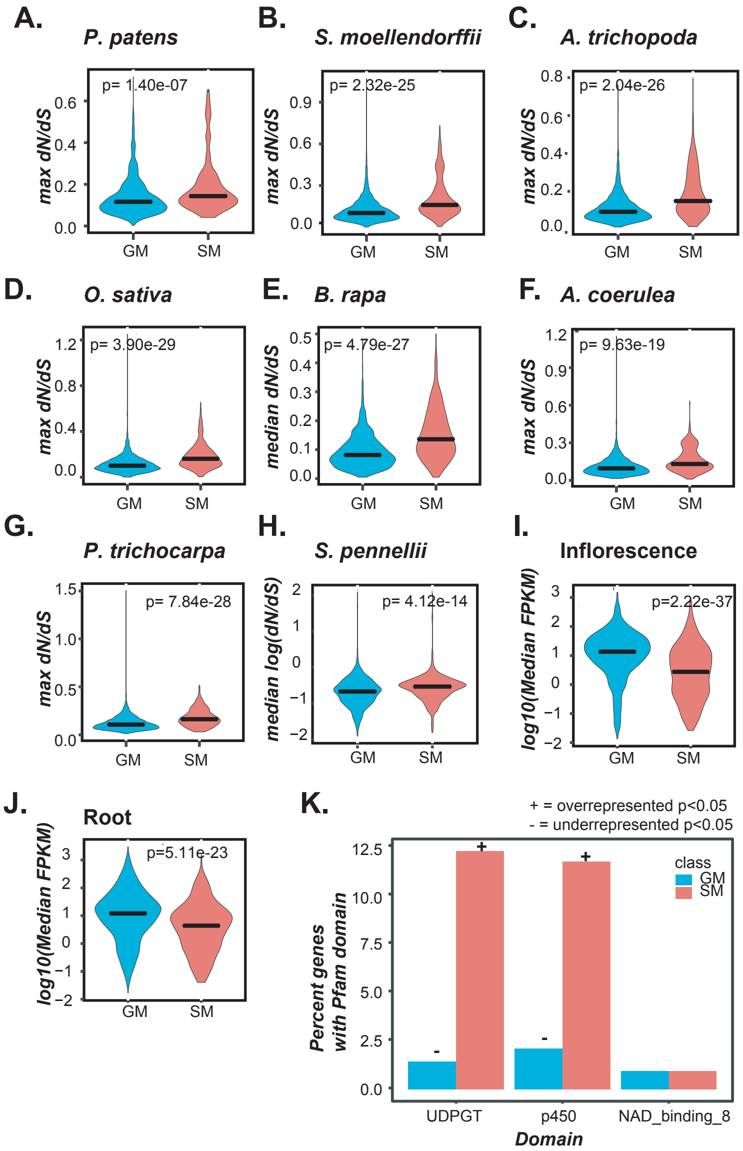

### Supplemental Figure 3

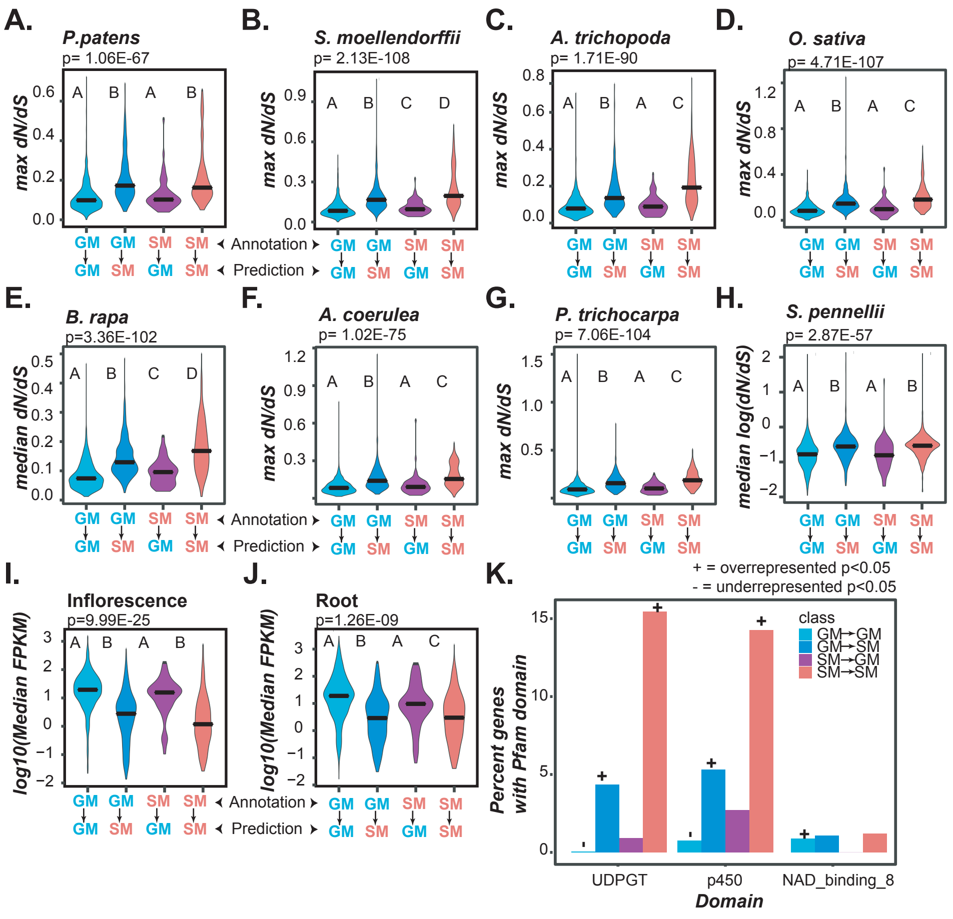

#### Supplemental Figure 4

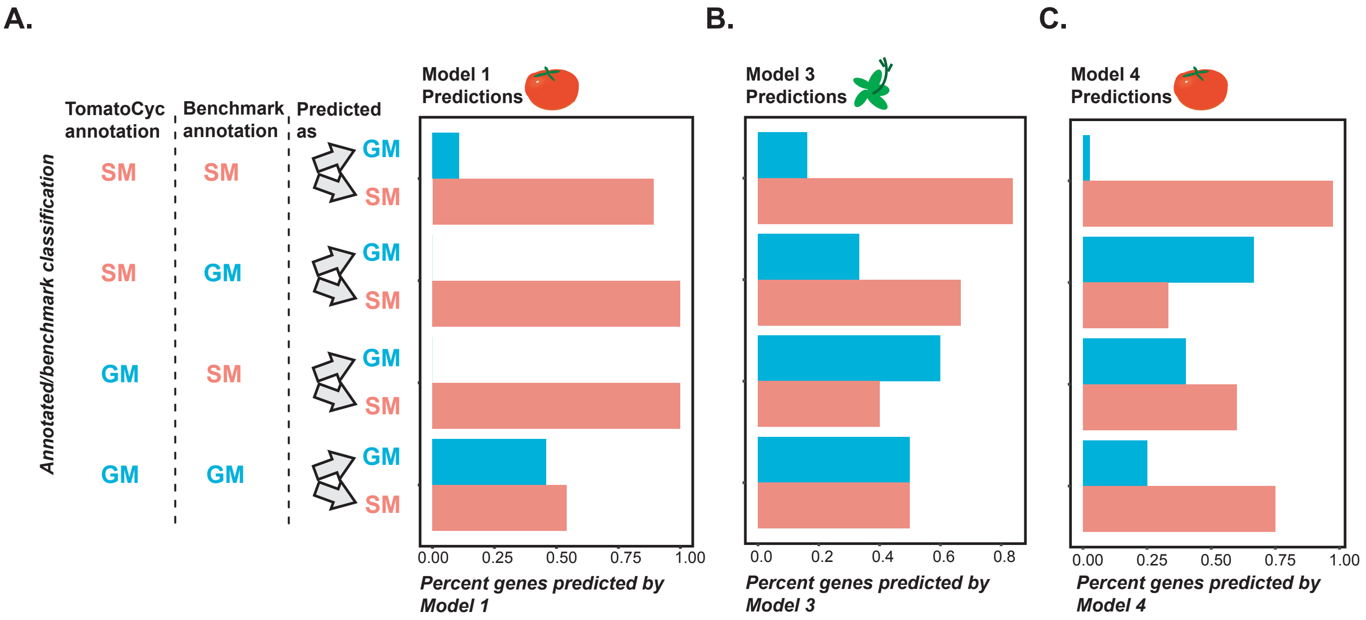

#### Supplemental Figure 5

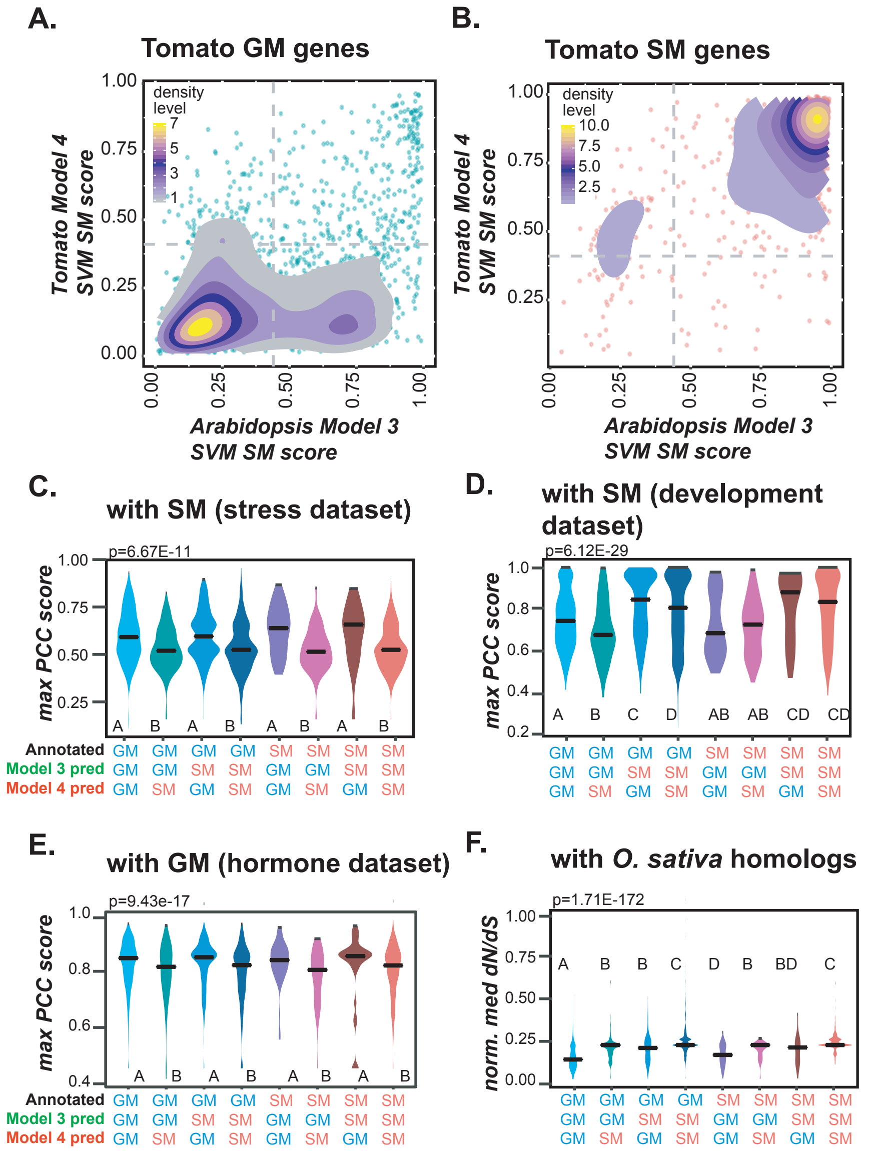

### Supplemental Figure 6

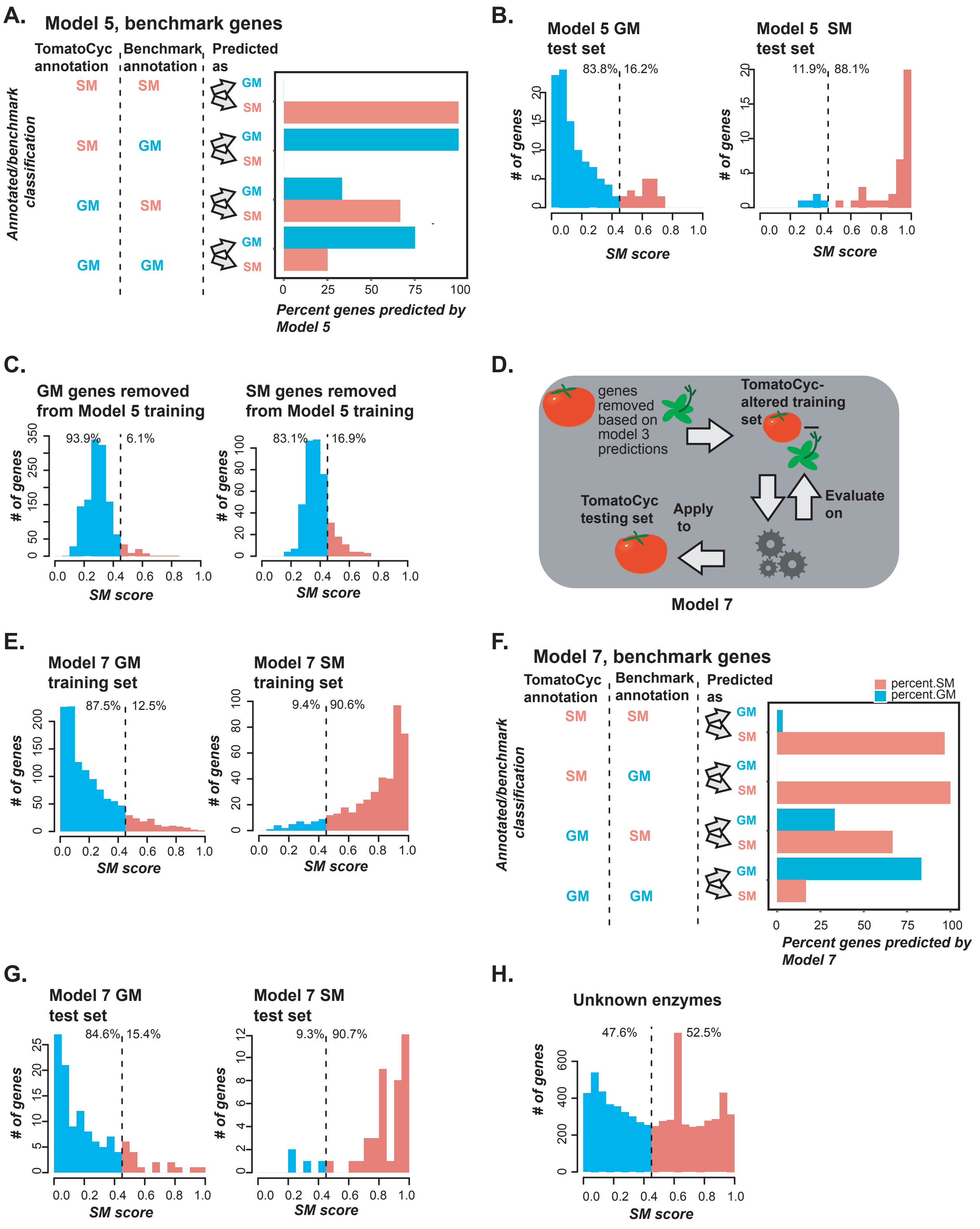

**Supplemental Figure 7**

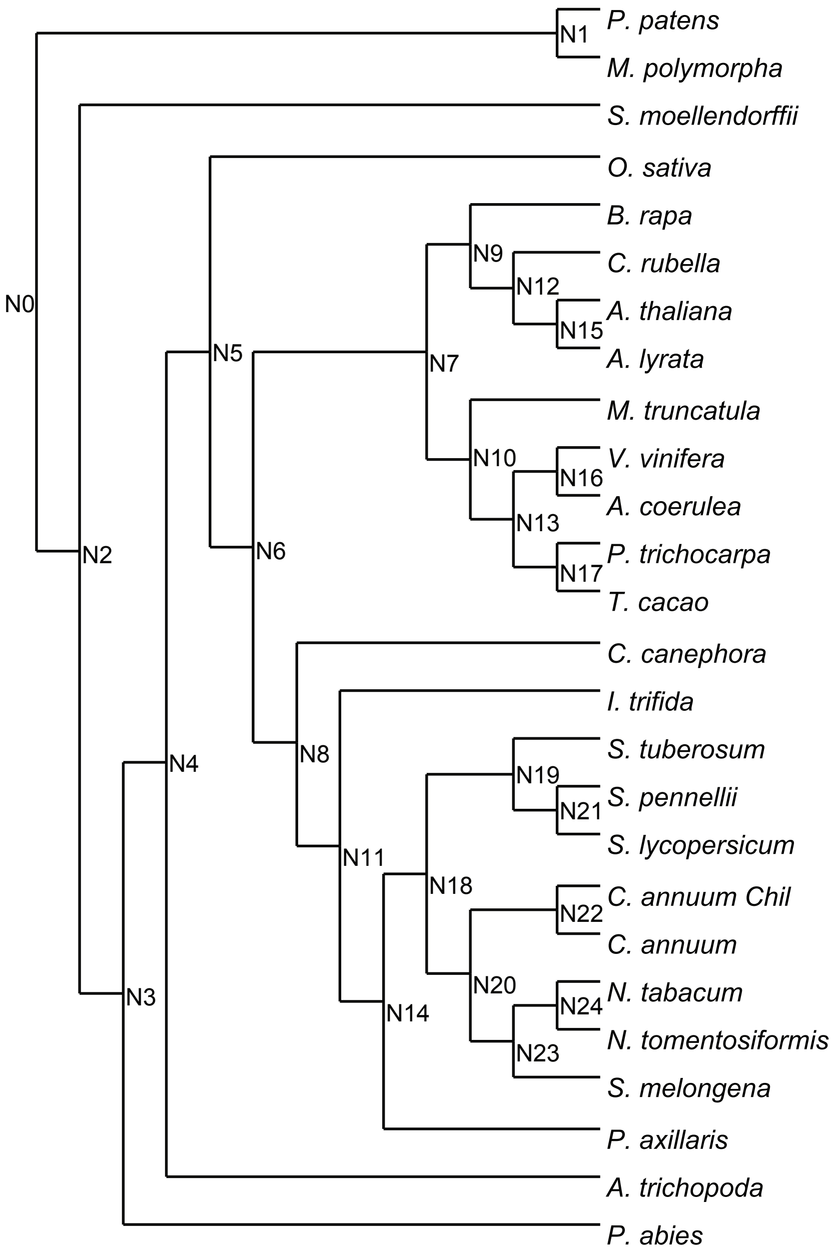
