## Supplementary material for "Within and cross species predictions of plant specialized metabolism genes using transfer learning": Dataset S1: Dataset_S1_features_legend.txt

Dataset S1: Full feature Dataset				
feature category	description of features in category	continuous or binary	number of features	key
evolutionary	"maximum or median dN/dS value of tomato gene to its homologs in various species, whether tomato gene has a homolog in a given species"	both	78	"dN/dS values are labeled as follows:
Slycopersicum_v_species_median/maximum dN/dS
Homologs: labeled as species.csv; value is for presence/absence of homolog"
duplication	"duplication type (tandem, syntenic, proximal, transposed), duplication node (approximation of when gene has last duplicated, corresponds to nodes in species tree), counts of gene neighbors, counts of gene paralogs (gene family size), pseudogenes (whether gene has pseudogene or not)"	both	26	"Duplication Features:
NeighborsXXCount: count of number of neighboring genes of XX type
XX.pairs: presence/ absence of this type of duplicate for given gene
node_c_XX: corresponds to duplication node on species tree where 0 is most ancient and 24 is most recent. See tree (Figure S7).
node_single: whether gene has no duplicate (1)"
expression	"number of times a gene is up or down regulated under certain conditions (breadth), median, max, or variation (MAD) under different data sets"	continuous	280	"expression breadth: XX_up or dwn: feature where XX represents dataset (see Table S7 for dataset description) and counts the number of conditions a gene is up-regulated (up) or down-regulated (dwn) in this dataset. groups of datasets include hormone, mutation, development, circadien, biotic and abiotic stress. XX_MAD: MAD is the mean absolute deviation measure for that gene (variance measure) under XX dataset
XX_med: median value of gene for XX dataset
XX_max: maximum value of gene for XX dataset
_FC : fold change values
_FPKM: FPKM values."
co-expression	"correlation values to SM, GM, or DF genes under different data sets"	continuous	1476	"correlation measures: labeled as: algorithm_FC or FPKM_dataset_SM/GM/DA_maximum or median
algorithms include: Spearman�s rank, Pearson�s Correlation Coeifficient (PCC), corpcor (Partial correlation).
dataset descriptions can be found in Table S7."
co-expression modules	presence/ absence of a gene in a co-expressed module under different data sets	binary	1194	"For clustered modules:
labeled as: algorithm_FC or FPKM_dataset_K-value_c_cluster number
algorithms are: cmeans, kmeans, hclust (hierarchal clustering) which has 3 types: average, ward, and complete; and WGCNA (weighted gene co-expression network average)
dataset descriptions can be found in Table S7."
domains	presence/ absence of Pfam domain in a gene	binary	4232	"Each feature is a Pfam protein domain and 1,0 determines presence,absence of domain in gene"
total number of features:			7286	
